# Multicellularity and virulence in the budding yeast, *Saccharomyces cerevisiae*

**DOI:** 10.64898/2026.10.26.761754

**Authors:** Guodan Zhong, Luke I. Ekdahl, Armaan Salchak, Atari Abundo, Mariana Vanpelt Chan, Grace Bellamy, Helen Murphy

**Author notes:** To whom correspondence should be addressed: Corresponding Author: Helen Murphy, Department of Biology, William & Mary, Integrated Sciences Center, Room 3035, 540 Landrum Drive, Williamsburg, VA 23185.

## Abstract

In the real world, microbes must cope with environmental fluctuations, compete for scarce resources, and avoid predation. One adaptation to survive this tangled web is the formation of multicellular aggregates, such as biofilms, which are known to be associated with pathogenicity. Despite the importance of the yeast *Saccharomyces cerevisiae* to biomedical research, little is known about the natural variation in its multicellular, biofilm-related traits. We investigated five multicellular phenotypes in a diverse, worldwide panel of ∼1000 isolates. We found that all traits varied significantly and ∼2/3 of the isolates expressed at least one multicellular trait. Both recent ecology and evolutionary history explained some of this variation, with clinical isolates exhibiting increased levels of three biofilm-related traits. When tested against wax moth larvae, hyper-multicellular isolates tended to be more virulent than non-multicellular isolates. Genome-wide association analysis on each trait suggested a genetic architecture of many small effect loci, compatible with polygenic adaptation. Only four genes, *SLN1*, *ECM5*, *MLP2*, and *ECM9*, were shared among the traits; deletion of *MLP2* and *ECM9* verified their association with multicellularity. Our results show that this yeast expresses far more multicellularity than previously appreciated and suggest that the aggregative phenotypes are likely evolving independently.

## INTRODUCTION

Most microbes live in spatially structured communities in which they interact with cells from their own and other species. Within this context, many have evolved the ability to act as a collective, forming a variety of multicellular structures such as swarms, slugs, fruiting bodies, and biofilms^1,2^. These phenotypes allow microbes to work together to forage, reproduce, and protect the community from environmental stressors. When found in medical settings, some multicellular phenotypes, particularly biofilms, have been associated with pathogenicity and virulence, as well as increased resistance to antimicrobial compounds^3–6^. While bacterial biofilms have been studied extensively, less attention has been given to those of fungi^7^. Opportunistic fungal infections are on the rise^8,9^, and many of the high-concern species are environmental fungi^10^. Understanding how traits that may be related to pathogenicity exist in the environment is an important complement to studying fungal pathogenesis.

Here, we investigate the multicellular phenotypes of the budding yeast, *Saccharomyces cerevisiae*, which is capable of growing in a variety of aggregative, filamentous forms^11,12^. *S. cerevisiae* is known for its fermentative properties and serves as a major biomedical model organism, but it is also found around the world in a wide range of ecological niches, from temperate forests to vineyards to cactus fields, and can act as an opportunistic pathogen of immunocompromised individuals^13–17^.

This diverse, worldwide population contains a significant amount of genetic and phenotypic variation^18,19^. We take advantage of a global panel of ∼1,000 isolates to investigate variation in the multicellular traits: their prevalence, distribution, ecological associations, and correlations between them. In a subset of twenty strains, we further test whether multicellularity is indeed associated with virulence. Finally, we use our phenotype data in combination with published genotype data to understand the natural genetic variation that exists for the different traits.

Our research centered on five multicellular traits, three of which are induced on solid medium. First, colonies that exhibit complex colony morphology (CCM) have hallmarks of fungal biofilms^3^: an extracellular matrix protecting cells and facilitating nutrient flow^20,21^; specialized cells and growth zones; expression of drug efflux pumps; and adhesins attaching cells to one another^22^. CCM is induced through carbon limitation. Our experiments limited the two different carbon sources most likely encountered by yeast in the environment and that have been shown to induce CCM: glucose and ethanol^23,24^. The second phenotype is pseudohyphal growth (PSH). Pseudohyphae, which are chains of attached, elongated cells that spread over a surface and are presumed to represent a foraging strategy, are induced in nitrogen-limiting conditions^25,26^. The closely-related third phenotype is invasive growth (IG)^27^, which is sometimes used as a proxy for PSH^28,29^. Pseudohyphae can grow into the agar surface. When surface growth is washed away, the pseudohyphae leave behind a visible footprint. The remaining two multicellular phenotypes, flors and plastic adherence, are expressed in liquid medium. The formation of flors or velum, which are floating mats of attached cells in an extracellular matrix^30^, are most commonly found in sherry and wine making^31^. The final phenotype, adherence to a plastic surface (PA), occurs as layers of surface-attached cells and represents a general adherence phenotype. As the amount of microplastics in the environment increases exponentially^32,33^, PA may have important ecological implications. Environmental isolates vary in their ability to express these phenotypes^23,34–36^, and previous research suggests the traits are not necessarily correlated^35,37^.

Environmental selection may favor these five phenotypes individually or multicellularity as a general trait. Invasive and pseudohyphal growth have been found to be associated with virulence in *S. cerevisiae*^38–40^, as have, more generally, biofilms in pathogenic fungi^41^; however, it is unknown which, or how, each of these lab phenotypes represents a growth form important for survival and expansion in the natural environment. Moreover, genetic studies of these traits have mostly focused on lab strains.

The genetic basis of CCM, PSH, and IG have been examined in the Σ1278b and S288c lab backgrounds. Published studies have used deletion genetic screens, genome-wide approaches, and forward genetics, ^28,29,37,42–50^ and have found hundreds of genes and multiple evolutionarily-conserved pathways contributing to the regulation of their expression. The pathways include nutrient-sensing ones, such as the filamentous MAPK (fMAPK), cAMP-PKA, TOR (target of rapamycin), SNF1, as well as those that sense other environmental conditions, such as the pH-sensing Rim101. These pathways can influence each other through interactions of their major protein kinases^51^ and through co-regulation of downstream targets, such as *FLO11*, the main cellular adhesin responsible for cell-cell and cell-surface attachment^52^. Genome-wide investigations into PA^53,54^ and genetic studies investigating flor formation^55^ have confirmed the central role of Flo11 in multicellularity. *FLO11* has the largest promoter in the yeast genome^56^ and serves as a hub for transcription factors from multiple signaling pathways^57^, as well as chromatin remodeling activity^58^ and contains two non-coding RNAs whose expression contributes to *FLO11* regulation^59^. While the majority of this research has been in a limited number of lab backgrounds, some studies have examined a subset of multicellular traits using a handful of clinical^46,50^ and environmental isolates^49,60–63^.

The known genetic basis of these phenotypes suggests many potential targets for allelic variation to accumulate and modulate multicellularity in natural populations. When considering the possible types of variants segregating globally, the complexity and interconnected nature of the signaling networks allows for the possibility of major effect variants that influence multiple traits. On the other hand, since so many genes and pathways are involved, and based on the observation in a lab background that hundreds of genes were privately associated with each trait, it may be that each phenotype is modulated by its own set of small effect allelic variants. Such variants, associated with complex, continuous traits like multicellularity, are often invisible in forward genetics as they cannot produce observable biological when tested individually in the lab.

While other studies have found variation in filamentous traits in clinical^64^, vineyard^30,34^, and smaller environmental^35^ strain panels, *S. cerevisiae* is still largely considered a simple, single-celled yeast. The extent to which the many possible multicellular phenotypes exist globally is unknown. Furthermore, natural genetic variation in multicellularity has not been studied using a population-level approach. Here, we use high throughput phenotyping, automated image analysis, and genome-wide association studies to investigate the phenotypic and genetic variation in multicellular traits in the 1000 genomes collection. Taking this global view can reveal whether specific phenotypes or genetic variants are associated with particular environmental conditions or evolutionary lineages, and whether a phenotype has evolved independently multiple times.

## RESULTS

### Multicellular Phenotypes

We first determined the prevalence and distribution of the phenotypes among the 1000 isolates (Figure 1), and included only one of the complex colony phenotypes (low dextrose, LD) to avoid bias. Surprisingly, ∼2/3 of the strains expressed at least one multicellular phenotype, with 1/4 expressing 3 or more (Fig 1A). Each individual phenotype was expressed in at least 20% of the isolates, with PSH being the most prevalent at 40% (Fig 1B-black points). Within each phenotype, there was a wide-range of expression (Fig 1C), where extreme levels of multicellularity were observed. PSH can have enhanced expression when isolates are on the edge of the agar plate. Thus, the observed distribution of PSH was influenced by edge effects, while the presence/absence data were not. Overall, the phenotypic results suggest the importance of collective behavior in the well-known budding yeast, as demonstrated by its high frequency in these environmental isolates. A collection of sample images of each phenotype for each isolate are available at https://y1000multicellularitymurphy.wm.edu, and analysis of the data for complex colonies induced by YPD and low ethanol can be found in the Supplementary Material.

**Figure 1:**
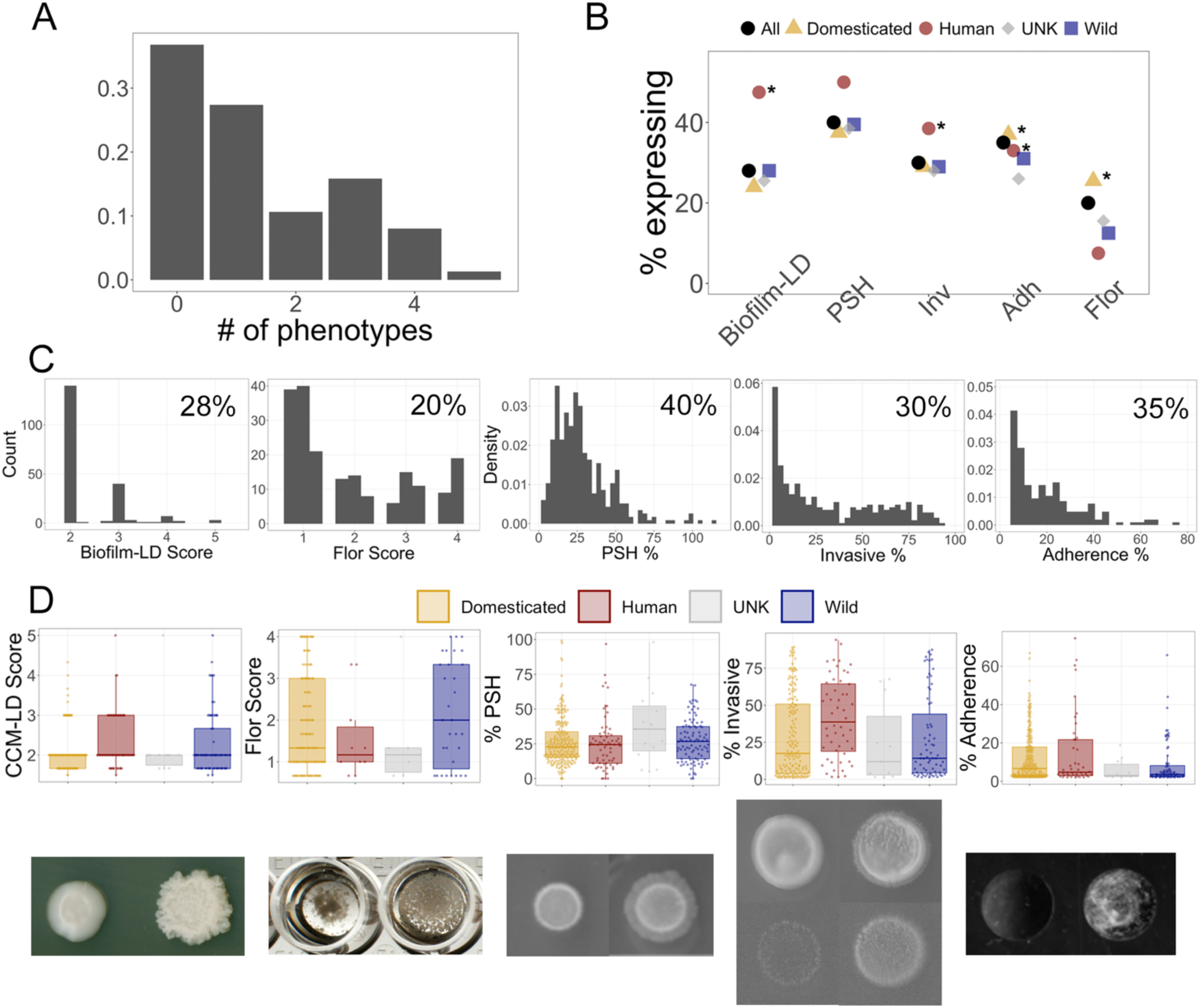
Prevalence and distribution of multicellular phenotypes in the 1000 genomes collection. **(A)** Proportion of isolates in the collection that exhibited different numbers of the following multicellular phenotypes: CCM-LD, flor, PSH, IG, and PA. Complex colonies on low dextrose represented biofilms; strains had to score at least 1.67 to be included (i.e., at least two of the three replicates were scored at 2). Similarly, isolates had to score 0.67 for flors, >1% for IG, and >2% for PA. Strains with PSH were computationally sorted (see Methods). **(B)** Proportion of isolates expressing each multicellular phenotype. The number of isolates in each category is as follows: human (135); domesticated (579); wild (245); unknown (39). The data for each phenotype were analyzed using an OLS (see Methods) with “wild” set as the reference category. Ecotype categories with significant coefficients are denoted with an asterisk. **(C)** Phenotypic distributions for the subset of isolates that exhibit the phenotype (i.e., excluding the non-multicellular isolates). The inset lists the percentage of isolates in the panel that exhibit the phenotype and on which the distribution is based. **(D)** Average phenotype values for all of the isolates in each ecotype category. Bottom row: Sample images that demonstrate the extremes of the phenotypes.

The relationship between multicellularity and different features of the isolates was investigated. This panel can be divided in at least two ways which encompass different ecological and evolutionary attributes of the isolates. The first is the broad ecology of the isolation substrate (ecotype), which may not directly reflect long-term evolutionary history. This categorization tells us where the isolate was found and represents our best estimation of the most recent environmental pressures the isolates experienced. The second possible division is phylogenetic clade. This categorization represents the long-term evolutionary history and genetic similarity among isolates, but does not explicitly account for information about the recent ecology of the strain.

First, the relationship between multicellularity and ecotype was investigated (Tables S1, S2). The prevalence and distribution of multicellularity in each isolation ecotype can be seen in Fig 1B (colored points) and Fig 1D. The human ecotype exhibits higher levels of CCM and invasive growth, suggesting more extreme versions of these phenotypes in the isolates in which they are expressed. The data were analyzed using an ordinary least-squares regression (OLS) framework setting “wild” as the reference category, as *S. cerevisiae* likely originated in a forest habitat^65^. With the exception of PSH, isolate ecotype explained a statistically significant, but small amount of the variation in the phenotypes (∼1% - 3%). This limited explanatory power is not surprising since there are only four categories and the analysis ignores the evolutionary history of these highly diverged isolates. Rather, it is noteworthy that this categorization explained any variation at all and that human-derived isolates were found to exhibit greater expression in invasive growth and in biofilm formation on both glucose- and ethanol-limited medium, which supports the association of these phenotypes with clinical relevance. In contrast, and as expected, domesticated isolates showed elevated flor formation. Interestingly, plastic adherence, which is a general adherence phenotype, was elevated in both human and domesticated categories. Taken together these results support the observation that human isolates express elevated levels of certain forms of multicellularity.

We next investigated the relationship of multicellularity with phylogenetic clade. Using OLS regression with a large “mixed origin” clade as the reference, the analysis explained 14-25% of the variation in the phenotypes. The increase in explanatory power is not surprising, as the data are divided into 34 categories based on their genetic similarity (clades and subclades). The results suggest that certain clades have a greater tendency toward some multicellular phenotypes and that particular clades, such as “French Dairy”, “African Beer”, and “French Guiana-Human,” exhibit elevated levels of multiple phenotypes (Fig 2, Table S3). These results highlight the importance of evolutionary history in multicellularity.

**Figure 2:**
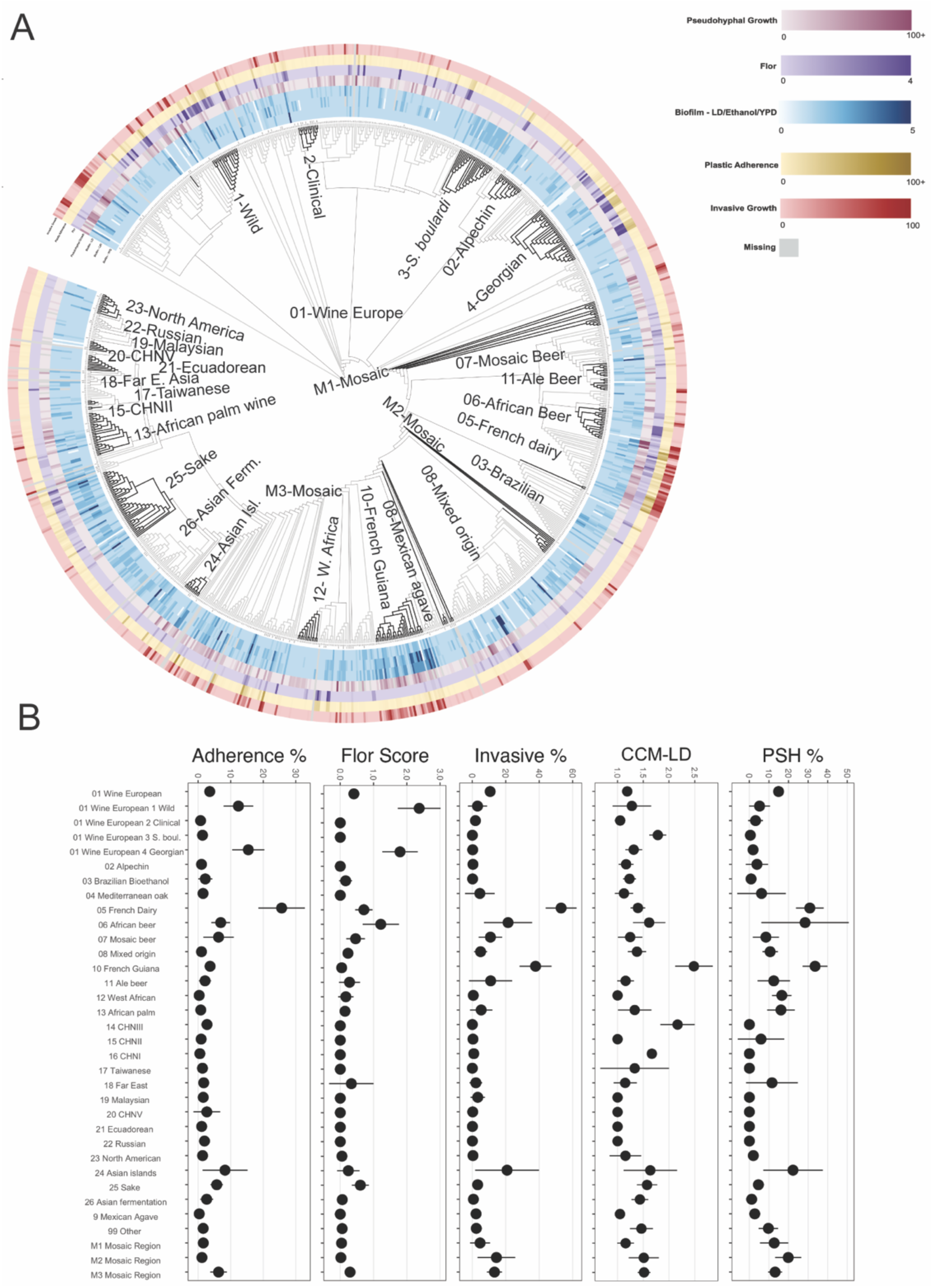
Multicellular phenotype expression in phylogenetic clades. **(A)** Cladogram of the 1011 genomes panel with phenotypes. **(B)** The average expression of all the isolates in a phylogenetic clade for each of the five multicellular phenotypes (± 2 sem). Complex colony formation on low dextrose represents biofilm formation.

To quantify the extent to which trait similarity reflects shared evolutionary history, we calculated the phylogenetic signal using Pagel’s Lambda^66,67^. Lambda ranges from 0 to 1, where a value of 0 indicates that trait variation is independent of phylogeny (no signal), and a value of 1 suggests that trait variation is strongly correlated with shared ancestry. There is a very weak phylogenetic signal across all multicellular phenotypes (0 < λ < 0.1, *p* < 0.05; Table 1).

**Table 1:** Extent of shared evolutionary history on trait values.

| Phenotype | $\lambda$ | *P*-value |
| --- | --- | --- |
| Plastic adherence | 0.079 | 0.004 |
| Invasive growth | 0.033 | 0.012 |
| Pseudohyphal growth | 0.041 | 0.006 |
| Biofilm colony (LD) | 0.038 | 0.011 |
| Biofilm colony (LEt) | 0.030 | 0.02 |
| Flor | 0.042 | 0.013 |

We next sought to determine the pairwise association between the multicellular phenotypes in an evolutionary context using phylogenetically independent correlations (Figure 3A; Table S4). This analysis seeks to determine if different combinations of phenotypes co-occur among the isolates, independent of shared of evolutionary history. Not surprisingly, the strongest correlation was between CCM induced on two different limiting carbon sources (glucose and ethanol; *r* = 0.52). The next strongest correlation was between IG and PSH (*r* = 0.49). Finally, the third strongest correlations were between IG and CCM on ethanol (*r* = 0.41), and flor formation and PA (*r* = 0.41). Thus, it appears that when multicellular phenotypes arise, they are more likely to occur with certain other phenotypes.

**Figure 3:**
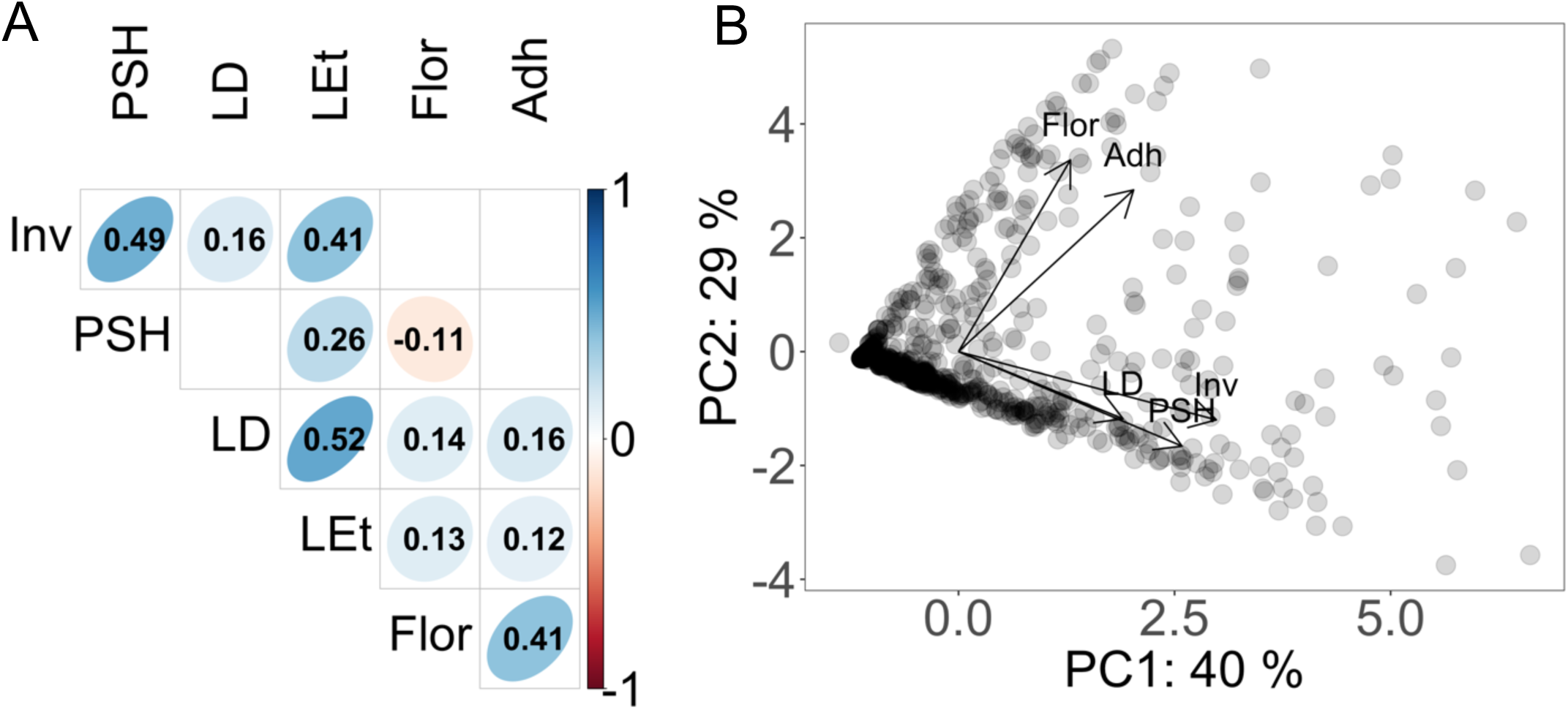
Relationships between the phenotypes. **(A)** Pearson’s correlation between pic-corrected values for the phenotypes. Only statistically significant correlations are shown. **(B)** Principal Component Analysis (PCA) of the five multicellular phenotypes, with CCM in LD representing the biofilm phenotype. PC1 (39.8%): 0.259*Flor + 0.381*LD + 0.597*Inv + 0.517*PSH + 0.405*Adh. PC2 (29.2%): 0.674*Flor - 0.236*LD - 0.240*Inv - 0.332*PSH + 0.568*Adh.

Finally, the data were combined into a principal component analysis (PCA) to investigate the relationship between all the phenotypes and determine whether this environmental panel contained naturally-occurring “hyper-multicellular” isolates similar to those that evolved in a laboratory experiment^68^ (Figure 3B). Most isolates cluster in one of two groups: those expressing flors, PA, or both, and those expressing some combination of CCM, IG and PSH. Only a small subset of isolates exists in the phenotypic space that strongly expresses both suites of traits. However, such isolates do exist in this panel. Interestingly, when a PCA is performed on phylogenetically independent contrasts, the phenotypes in the second suite of traits (CCM, IG, PSH) appear to be more independent of one another, and no hyper-multicellularity is observed (Figure S2). Thus, the phenotypic relationships and hyper-multicellularity observed in the panel are strongly influenced by evolutionary history.

### Virulence

In a previous experiment in which hyper-multicellular strains evolved through selection on plastic adherence, the strains were also incidentally more virulent^68^. To test whether hyper-multicellularity is associated with increased virulence more generally, we used our data to identify naturally-occurring strains that displayed hyper-multicellularity across all traits and strains that exhibited no multicellularity, designated “high” and “low” multicellular, respectively. We also identified strains that exhibited individual/correlated multicellular traits, PSH/IG and flor/PA; CCM was always associated with multiple other traits, so not considered alone (Table S5). The strains were injected in the larvae of the wax moth, *Galleria mellonella*, and larval survival was monitored over a one-week period. As was observed in the evolution experiment, the hyper-multicellular strains tended to be more virulent (Figure 4). When wax moth larvae survival was modeled with multicellular status as a fixed effect and “low” set as the reference category, and with strains included as a random effect, the “high” category showed increased virulence (HR = 1.45, *p* = 0.0616), while PSH/IG (HR = 1.10, *p* = 0.6388) and Flor/PA (HR = 1.09, *p*=0.6816) did not (Table S6). Still, there was a lot of variation among strains that was not explained by multicellular status (Figure 4B; Table S7). Virulence is clearly a multifaceted trait and multicellularity is not sufficient to explain it; however, it does appear that multicellularity can be a contributing factor. While the statistical significance is marginal, it is notable that strains with very different genetic backgrounds and evolutionary histories, and chosen only because of their multicellular status, still show a signal of increased virulence with hyper-multicellularity. These findings suggest that multicellularity may be a contributing factor to the multifaceted trait virulence.

**Figure 4:**
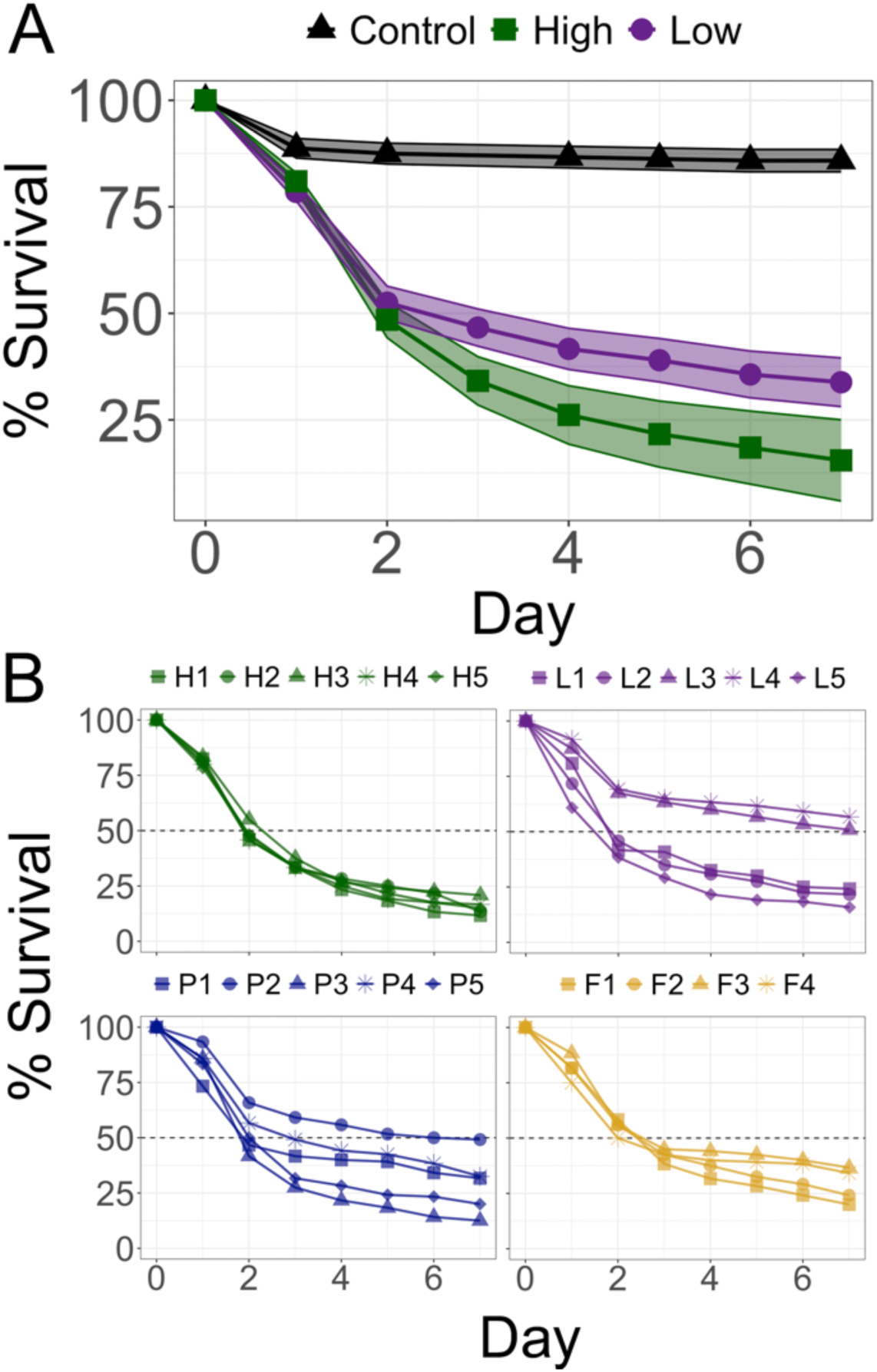
Hyper-multicellularity and virulence. *G. mellonella* survival curves for isolates with low multicellularity, PSH/IG, flor/PA, or hyper-multicellularity. Each strain was injected into 120 larvae; points represent Kaplan-Meier estimates. **(A)** Survival curves with confidence limits for low- and hyper-multicellular isolates; black triangles represents the control treatment injected with sterile water. **(B)** Survival for individual strains in each category.

### Genetic Basis of Individual Phenotypes

Previous research suggests that each of the multicellular phenotypes has its own complex genetic basis^37^ with some notable overlap, including in certain signaling pathways and in cellular adhesins, such as Flo11. In order to determine the type and amount of allelic variation in this diverse world-wide panel, we conducted a genome-wide association analysis (GWAS) on each phenotype using available genome sequence data^18^ and genotype files^69^ (Figure 5). This allowed us to identify genetic variants associated with phenotypic variation without relying on predefined candidate genes.

**Figure 5:**
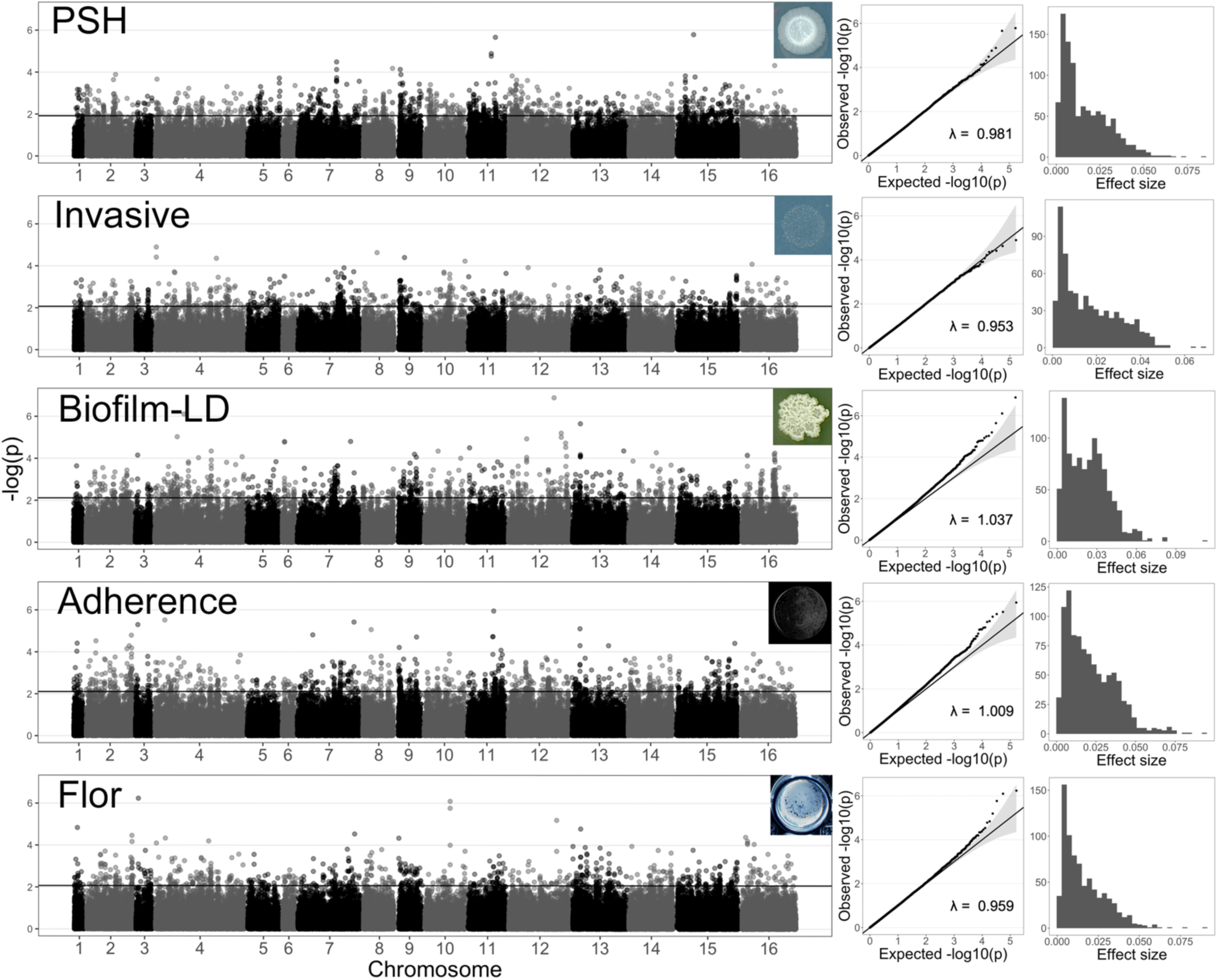
Genome-wide analysis of the multicellular phenotypes. Each row represents the analysis of a single phenotype. The first column contains the Manhattan plot, which plots the -log_10_(*p*-values) for the nuclear genomic SNPs; mitochondrial data not included. The second column contains Q-Q plots, which plot the observed *p*-values and the expected *p*-values under a null distribution; each contains a genomic control inflation factor (λ). The third column contains effect size distributions for the statistically significant SNPs. After normalization, the range of phenotypic values were: PSH -5.2 – 5.2, IG -5.2 – 5.2, CCM-LD -3.9 – 3.4, PA -5.2 – 5.2, flor -0.6 – 2.0.

Once again, the analysis included a single biofilm phenotype (CCM on LD) to avoid skewing the results. All genomic control inflation factor (λ_GC_) values are close to 1, suggesting that the population structure was correctly stratified for all GWAS. The deviations between the expected and observed -log_10_(p) values in quantile-quantile plots suggest that the genetic variants in the populations are only weakly associated with PSH and IG, while for CCM, flor, and PA, there are likely strongly associated variants (Figure 5). With this caveat in mind, we further analyzed the results.

The heritability of the traits ranged from 0.40-0.67 (Table 2). Each phenotype had its own unique set of variants, with limited overlap among them, confirming the independent, complex genetic basis of different forms of multicellularity. Combined, there were 4157 significant loci detected across the five phenotypes; these translated into 3499 unique SNPs in 1640 genes and 508 intergenic variants (Tables 2 and S8). There were very few variants with moderate effect sizes; most variant effects were estimated to be small (Figure 5). As the phenotypic data were transformed to meet the assumptions of the GWAS, the range of the final transformed values was as low ∼2.5 for flor formation and as high as ∼10 for PA, PSH and IG. Thus, an effect of 0.01 on the phenotype is indeed quite small. Of the twenty largest effect variants for each of the phenotypes, the majority were private and not shared: CCM-LD (15/20), invasive growth (15/20), PSH (19/20), PA (12/20), and flor (10/20). Thus, it appears that most natural variation only very slightly modulates multicellularity, and the variation with larger phenotypic effects is generally specific to individual multicellular traits. When looking at the overlap among genes associated with each phenotype, the pattern of correlated traits is recapitulated in the genetic overlap. For comparisons with flor, the flor-IG comparison has the most overlap in genes, while for comparisons with PSH, the PSH-IG has the most overlap (Figure 6C; Table S9).

**Figure 6:**
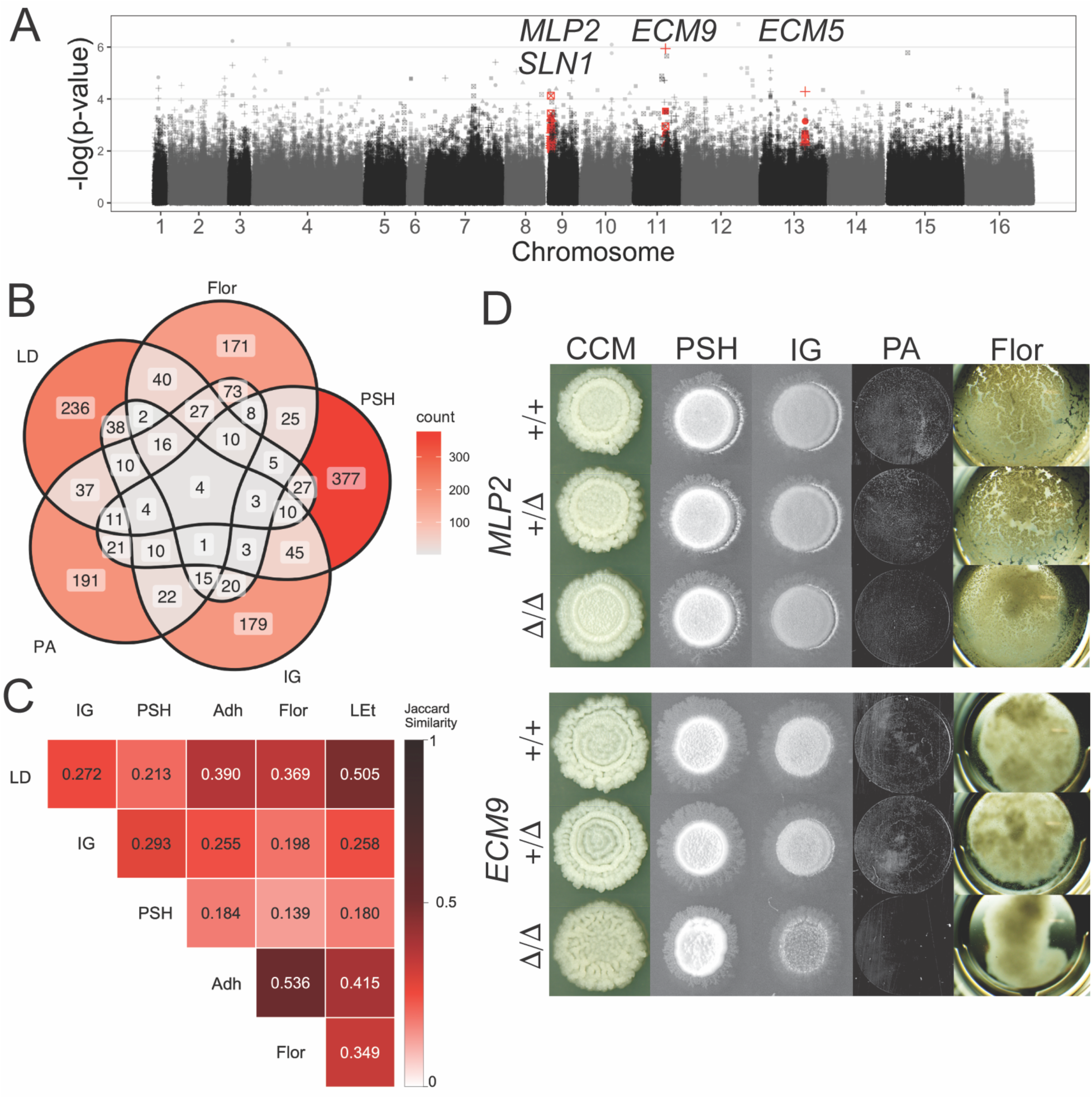
Overlap in the genetic basis of multicellular phenotypes. **(A)** Manhattan plot that includes all five phenotypes, with the four genes significant in all of them highlighted in red; each phenotype is represented by a different point style. **(B)** Venn diagram of genes containing significant SNPs, although the individual SNPs within a gene may differ between phenotypes. **(C)** Shared associations were quantified using gene–phenotype relationships from Table S9. Pairwise overlap was quantified using Jaccard similarity: J = S / (N1 + N2 - S), where S is the number of shared genes and N1 and N2 are the numbers of genes associated with the two phenotypes, respectively. **(D)** The five multicellular phenotypes in wildtype, heterozygous deletion, and homozygous deletion strains for *MLP2* and *ECM9* in clinical isolates. Phenotypes based on 8 days, instead of 5.

**Table 2:** Summary of the GWAS results. Loci represent all statistically significant variants from a linear mixed-model analysis in FaST-LMM, which also produced effect sizes; type of SNP was assigned using snpEff (see Methods). Some variants had multiple annotations. Narrow sense heritability (h^2^) was estimated using variants on nuclear chromosomes under the null model in FaST-LMM.

| Phenotype | Loci | Intergenic | Gene | Syn | Non-syn | Effect size (median) | Effect size (mean) | $h^2$ |
| --- | --- | --- | --- | --- | --- | --- | --- | --- |
| Flor | 695 | 98 | 422 | 286 | 173 | 0.0125 | 0.0165 | 0.575 |
| Plastic adherence | 836 | 84 | 459 | 431 | 172 | 0.0186 | 0.0217 | 0.552 |
| Biofilm colony (LD) | 933 | 101 | 479 | 473 | 187 | 0.0225 | 0.0233 | 0.467 |
| Invasive growth | 664 | 80 | 381 | 305 | 159 | 0.0138 | 0.0169 | 0.406 |
| Pseudohyphal growth | 1029 | 145 | 564 | 457 | 252 | 0.0117 | 0.0166 | 0.671 |

GO enrichment analysis revealed a total of 72 pathways that were significantly enriched among genes identified by GWAS for the various multicellular traits. The most strongly represented pathways included metabolic pathways, biosynthesis of secondary metabolites, cell cycle, MAPK signaling pathway, and meiosis, containing 244, 111, 56, 46, and 48 GWAS-associated genes, respectively. Together, these results indicate that variation in multicellular traits is associated with multiple fundamental cellular processes, suggesting a highly polygenic basis involving diverse biological functions.

### Genetic Basis of General Multicellularity

Investigation into the genetic overlap of the various traits uncovered only a single synonymous SNP shared among all of the phenotypes (in *ECM9*; Table S10). We therefore looked for overlap in the genes associated with the five multicellular traits, noting that different SNPs in the same gene may be associated with specific traits, but these multi-SNP containing genes may be targets for modulating multicellularity more broadly. There were four genes that were shared among all the phenotypes, and each had SNPs with small effect sizes (Fig 6A-B): *ECM9* (*YKR004C*), *MLP2* (*YIL149C*), *SLN1* (*YIL147C*), and *ECM5* (*YMR176W*)^70^.

First, *ECM9* is a subunit of geranylgeranyl transferase type III (GGT3) that transfers a geranylgeranyl to Ykt6, which itself is a conserved SNARE protein involved in membrane trafficking (specifically intra-Golgi and autophagic membrane fusion)^70,71^. Deletion mutants in the BY4741 background have fragile cell walls and decreased autophagy^71^. *ECM9* contained one variant common to all the phenotypes, and another one associated with invasive growth; the effect sizes ranged from 0.005-0.018. To verify the gene’s effect, it was deleted in an isolate that had the “high” version of the allele and exhibited elevated multicellularity, SACE_YDN. In the mutant, biofilm colony formation was altered, pseudohyphal and invasive growth decreased, and plastic adherence and flor formation were abolished, although there was only minor flor formation in the original strain.

Next, *MLP2*, is a myosin-like protein involved in spindle pole body organization and regulation of protein import into the nucleus during the mitotic cell cycle spindle assembly checkpoint^70,72,73^; 19 SNPs (10 non-synonymous, 9 synonymous) with effects ranging from 0.006-0.074, were associated with various combinations of the phenotypes. In order to verify *MLP2*’s effect on multicellularity, the gene was similarly deleted in an isolate with the high allele, SACE_YBJ. In this isolate, biofilm colony formation became less “rough”, and plastic adherence and flor formation decreased, but were not abolished. Pseudohyphal and invasive growth were unchanged.

Third, *SLN1*, is an osmo-sensing transmembrane histidine phosphotransfer kinase that functions as part of the HOG pathway^70,74^, a pathway known to be associated with CCM^42^. Three non-synonymous and two synonymous SNPs with effect sizes ranging from 0.007-0.024 were associated with various combinations of the phenotypes. Deletion mutants are inviable^74^; therefore the gene was not investigated further. Finally, *ECM5*, is a subunit of the Snt2C complex, which modifies chromatin in response to oxidative stress^70,75^; one non-synonymous and three synonymous SNPs, with effects ranging from 0.002-0.011, were associated with various combinations of phenotypes. Published data show that deletion mutants exhibit an enlarged size and large drooping buds with elongated bud necks^76^. Since there was already a known association with cell and bud morphology, this gene was not investigated further.

The subtle, but measurable effect of the *MLP2* and *ECM9* deletions on multiple multicellular phenotypes, and the identification of the genes *SLN1* and *ECM5*, with plausible roles in multicellularity, suggests the possibility that much of the allelic variation identified in our GWAS may be causal, but with such small effect sizes are unlikely detectable in the lab.

## DISCUSSION

In this study, we surveyed the multicellular phenotypes of a large, ecologically and geographically diverse collection of the biomedical model yeast, *Saccharomyces cerevisiae*. Previous research has shown significant growth rate^18^ and reproductive^77^ variation in the collection, but this kind of organismal variation has not yet been investigated. While previous studies have provided valuable insights into *S. cerevisiae* multicellularity, their focus on laboratory strains may have limited the diversity and complexity observed. Across the species panel, we found a substantial amount of variation in all traits measured, and a surprisingly large proportion of isolates expressing multicellularity. We observed some impressive biofilms and pseudohyphae, but most of the isolates that expressed multicellularity did not express an extreme version; instead, they expressed a moderate, yet detectable form. The phenotypes were being induced in the lab using conditions reported in the literature. However, there may be other conditions or stimuli that enhance multicellularity or even induce other forms of it. Recent work has shown nutrient conditions associated with plant matter may also induce filamentous growth^24^, although those conditions were not screened here. It is unknown what purpose these multicellular traits serve in the environment, and how they manifest when not in the lab. For example, complex colony morphology may serve as a good proxy for the ability to form a biofilm, but what a natural *S. cerevisiae* biofilm looks like and how it functions in the environment remains a mystery. Intriguingly, one study showed that biofilm expression could play a role in protection from predation by a macropredator ^44^, and another showed a competitive advantage to strains producing biofilms over those not producing them^78^. More studies are needed to elucidate the function of the different types of multicellularity in the environment. Still, our data suggest that the tendency to act collectively is widespread across the collection. Since microbes in the real world exist in spatially-structured and complex biotic environments, it may not be unexpected for this panel to exhibit the level of multicellular variation we observed. However, since much of the research that employs *S. cerevisiae* as a model uses lab strains, the morphological complexity observed in these natural isolates and the selective pressures favoring it are likely underappreciated.

Certain patterns in the phenotype data stand out. First, invasive growth and biofilm formation were elevated in human-derived isolates. IG and PSH have been reported to be associated with virulence in a limited number of clinical *S. cerevisiae* strains ^38–40^, and biofilm formation is known to be associated with pathogenicity in other fungi ^41^. Thus, our observations demonstrate the association of these phenotypes with clinical relevance in *S. cerevisiae* on a larger scale. In contrast, flor formation was associated with domesticated isolates. This supports the idea that fermentation settings select for traits that allow cells to adhere to one another, and sometimes form floating mats, a well-known phenotype associated with sherry production^31^. More broadly, our results show that patterns hypothesized to exist are in fact detectable in a large, ecologically diverse panel, even when not correcting for evolutionary history or genetic similarity. Interestingly, PA was associated with both clinical and domestic isolates, supporting the notion that it represents a general adherence phenotype. It is worth noting that as the abundance of microplastics increases exponentially^79^, there may be an entirely new ecological niche^80^ that could indirectly select for traits associated with pathogenicity and virulence^68,81^.

The ecotype analysis was complemented with a phylogenetic analysis. Certain clades showed elevated forms of multicellularity, but over the whole global population, the traits were only weakly associated with phylogeny (i.e., very small Pagel’s λ values). This suggests that both ecological niche (current selective pressures) and evolutionary history together partially generate the patterns we observe, explaining a bit less than half of the variation in the traits.

Similar ecological conditions have independently favored the evolution of similar multicellularity phenotypes across distinct evolutionary lineages. For example, despite the evolutionary distance between “French Dairy” and “Sake”, they exhibit similar levels of flor formation. Such phenotypic convergence might be explained by ecology, as most strains in these two clades were isolated from domestic environments. On the other hand, similar selection pressure may not be enough to explain our observations, as strains from the same clade isolated from the same ecotype have different multicellular profiles. For instance, AHR and Sace_YAD are strains in “Wine/European” isolated from clinical environment. While Sace_YAD shows a 32.3% mean pseudohyphal growth, AHR does not grow pseudohyphae at all. These observations suggest that ecotype and evolutionary clade do not fully explain variation in multicellularity, with finer-scale environmental and genetic differences also playing important roles.

Finally, while there were some correlations between phenotypes, expression of one phenotype did not necessarily predict the expression of others. This observation supports a previous report for a smaller collection of ∼30 isolates and their haploid derivatives, which found that various multicellular traits were only weakly or not at all correlated^35^. The correlations we observed clearly split the panel into two: those isolates expressing of PSH, IG, and/or CCM, and those expressing flor formation and/or PA. Notably, when evolutionary history was accounted for, the relationship in the first group disappeared (PSH, LD, IG). This result, combined with the observation that there are very few naturally occurring hyper-multicellular isolates, suggests that these phenotypes may be evolving independently of one another. Different selective pressures may be favoring each or some combination of the traits, but not necessarily an overall, generalized multicellularity.

Based on previous research, we hypothesized that the natural hyper-multicellular isolates would exhibit elevated virulence^68^. When tested, they caused elevated death rates in wax moth larvae compared to non-multicellular strains and strains that exhibited only one of the suites of multicellular phenotypes. Despite the lack of an abundance of hyper-multicellular isolates in the panel, and most likely in the environment, there is something about extreme multicellularity that leads to its association with virulence in this species. The isolates that were assayed had different genetic backgrounds, ecotypes, isolation substrates, and geographical origins. Thus, it is likely the case that it is the hyper-multicellularity itself that led to the increased virulence, but the selective pressure leading to this phenotype and the reason it is associated with virulence remains unknown.

Thus far, natural, segregating genetic variation in the multicellular traits has not been investigated on a large scale. When conducting GWAS using the 1011 genomes panel and correcting for population structure, major effect alleles can be detected with reasonable confidence, while alleles of small effect are more difficult to identify^82^, as are the effects of low-frequency variants^83^. As with all GWAS, spurious associations remain possible and are more likely in increasingly complex traits with small effect alleles. However, with this panel, GWAS has been successfully employed to uncover variants associated with life history traits^69^ and growth rate in various conditions^18^. Here, we used the panel to find variants associated with multicellularity.

Our results support a model that describes the genetic variation for each trait as many small effect loci with very few large/medium-effect variants. These results are not surprising when considering the number of genes shown to affect PSH, IG, and CCM in genetic studies on lab strains. It is worth noting that our analysis used published genotype files that did not include the accessory genes in the *S. cerevisiae* pangenome. These accessory genes have been shown to have a significant effect on transcriptional profiles^84^ and may be important in other phenotypes, such as the ones investigated here. This leaves open the possibility that there are major effect loci that we did not detect. More broadly, our results fit under the polygenic adaptation framework, where selection acts on multiple genetic loci associated with one trait^85^. Such adaptation does not require the same genetic changes across populations, where various genetic modifications in different populations can convergently achieve similar outcomes^86^.

Interestingly, there was very little overlap in the variants associated with each of the traits. This supports the idea that the traits are evolving independently from one another. When considering the overlap of genes (as opposed to specific SNPs), the correlated phenotypes showed more genes in common than non-correlated phenotypes. This observation makes sense in light of the evolutionarily conserved pathways underlying the control of the traits, although many of the genes uncovered in our study had not been implicated in any of the multicellular phenotypes before.

Only four genes were associated with all five of the multicellular phenotypes, and the estimated effect sizes of the SNPs were small. None had previously been associated with filamentous growth, although one, *SLN1*, is a known component of the HOG pathway, which has been implicated in CCM^42^, and deletion of another, *ECM5*, had been shown to cause abnormal cell morphology. The remaining two genes, *MLP2* and *ECM9*, were deleted in this study and both caused a decrease in multicellular traits. Since the deletions did not cause loss of the phenotypes, it is unlikely that the effect of individual SNPs would be detectable in the lab. However, the results suggest that despite the difficulty of detecting small effect loci and the possibility of spurious associations, at least some of the small effect loci we have uncovered are likely true associations with the phenotypes.

By studying the naturally occurring variants in this global collection, our data provides an important complement to studies that use deletion and overexpression collections in lab strains. In addition to determining whether a gene is associated with a trait, we are able to begin to understand which allelic variants may be responsible for creating the variation we see in environmental isolates.

Our phenotypic and genetic observations paint an emerging picture: the different multicellular traits in *S. cerevisiae* may be independent of one another. Given the many ecological niches and environments in which this yeast can exist, it is not surprising that different forms of collective behavior may be favored. It is possible that selection is favoring multicellularity for different purposes in different settings, although the details of the non-domestic environments and selective pressures remain largely unknown.

## MATERIALS AND METHODS

### Strains

This study investigated 998 strains of the 1,011 *Saccharomyces* genome panel^18^, provided by Paul Rowley (University of Idaho) in a 96-well plate format. The original three 384-well plates were divided into plates 1A-1D, 2A-2D, and 3A-3D, and included control strain FY4 in multiple wells on each plate. The clades associated with each strain were obtained from the original study, while the categorization of ecotype/ isolation origin were from a follow-up study^69^.

Two clinical isolates in the panel, originally from another collection^64^, were engineered to have gene deletions using a lithium acetate transformation^87^ supplemented with 3% DMSO and an antibiotic cassette (KanMX)^88^ targeted to the gene of interest. Primers containing 18-19 bp of homology to the cassette and 41 bp of homology to the gene were generated by PCR using Phusion enzyme (New England Biolabs) with the manufacturer’s recommendations. Colonies growing on YPD supplemented with G418 (200 ug/ml) were sporulated and dissected to isolate strains homozygous for the deletion. *MLP2* was deleted in SACE_YBJ/ YJM1190/ UM25A/ UCD-FST 08-191 to generate HMY918 (heterozygous for the deletion) and HMY 913 (homozygous for the deletion) using primers: MLP2-MX-for: 5’ ATGGAGGACAAAATATCTGAGTTTTTAAATGTTCCATTCGA *CGTACGCTGCAGGTCGAC* and MLP2-MX-rev: 5’ TGGAAGCTTGATCATTTGCTGGACTCTCTTTAACCTTTTTG *ATCGATGAATTCGAGCTCG*. *ECM9* was deleted in SACE_YDN/ YJM1083/ NRRL Y-10988 to generate HMY919 (heterozygous for the deletion) and HMY912 (homozygous for the deletion) using primers: ECM9-MX-for: 5’ ATGCAATCTTCTCTTCCCCTTTGCAGAGAATTTTTTGAAAA *CGTACGCTGCAGGTCGAC* and ECM9-MX-rev: 5’ CTAACTATCTGATTGTTCTTTCTTATTTTTATCCAAAAACT *ATCGATGAATTCGAGCTC*. Antibiotic inserts were verified with primers: MLP2-for: 5’ ATGGAGGACAAAATATCTGAGTTTTTA and MLP2-rev-5’ TGGAAGCTTGATCATTTGCTGGACTCT, and ECM9-for: 5’ ATGCAATCTTCTCTTCCCCTTTGCAGA and ECM9-rev: 5’ CTAACTATCTGATTGTTCTTTCTTATTTTTATCCAA.

### High-Throughput Phenotyping Assays

The phenotypes were investigated in three sequential assays, with each assay containing an entire set of A-D (one original 384-well plate), along with a control plate that contained 19 strains with known multicellular phenotypes (those used in the virulence experiments). The five plates (A-D, control) were assayed for all multicellular phenotypes at once, in replicates of three, on the same batch of medium. The five plates were cultured from the freezer in liquid YPD (1% yeast extract, 2% peptone, 2% dextrose) for 48 hours, then transferred to another plate for overnight growth using a 96-pin multi-blot replicator (V&P Scientific no. VP408FP6). Yeast cultures were then pinned to different media to induce various multicellular traits. With the exception of flor, all plates were scanned with an Epson Expression 11000 XL scanner with settings for black and white images at 1200 dpi. Each plate image was split into 96 individual images for computational analysis.

#### Biofilms/ Complex Colony Morphology (CCM)

The CCM phenotype was assayed using three different solid agar media in OmniTrays (Nunc 264728): (1) yeast-peptone-low-dextrose (YPLD) medium (1% yeast extract, 2% peptone, 2% agar, 0.5% dextrose), (2) yeast-peptone-low-ethanol (YP-LEthanol) (1% yeast extract, 2% peptone, 2% agar, 0.5% ethanol), and (3) YPD. The first two media created low carbon environments, while the third was a rich environment meant to act as a control. The CCM plates were imaged after 5 days of 30C incubation.

#### Pseudohyphal Growth (PSH)

PSH was screened on solid synthetic low-ammonium dextrose (SLAD) 2X medium (2% dextrose, 2% agar, 0.34% yeast nitrogen base without amino acids or ammonium sulfate, and 50 µl of 1M ammonium sulfate) in OmniTrays to create a nitrogen-limiting condition. The SLAD plates were made one day prior to pinning and imaged after 5 days of 30 C incubation.

#### Invasive Growth (IG)

The SLAD plates used to screen PSH were washed in order to analyze IG. After imaging PSH phenotypes, the SLAD plates were flooded with diH_2_O and gently washed with a gloved finger, then similarly rinsed, to remove the yeast colony remaining on the surface level of the agar. This “wash step” enabled the visualization of yeast cells penetrating the solid surface invasively. The washed SLAD plates were imaged. All liquid waste was collected for sterilization.

#### Plastic Adherence (PA)

PA was phenotyped in 96-well plates containing 200 ul liquid synthetic low dextrose (SLD) media (0.67% yeast nitrogen base without amino acids and with ammonium sulfate and 0.1% dextrose). After 2 days of 30C incubation, plates were emptied, then filled with diH_2_O, emptied, and blotted while inverted on an absorbent surface; the wash step was performed 3 times. The plate was dried in a 30C incubator and the remaining adherent cells were imaged. All liquid and solid waste was collected for sterilization.

#### Floating Mats (Flor)

Flor was phenotyped in 96-well plates containing 200 ul of liquid synthetic dextrose (SD) medium (0.67% yeast nitrogen base without amino acids and with ammonium sulfate and 2% dextrose) per well. Flor was scored under a light microscope after 5-day 30C incubation manually using the following scale (Figure SX): 0-no floating cells, 1-scattered floating cells that cover a small area of the liquid surface, 2-scattered floating cells that cover a large area of the liquid surface, 3-floating cell clumps that cover some area of the liquid surface, 4-floating cell clumps that cover all liquid surface.

### Computational Image Analysis

#### CCM

We used a 5-point scale to quantify the complexity of the biofilms. A deep learning pipeline implemented with TensorFlow and Keras packages, two widely used open-source libraries for machine learning (ML) and artificial intelligence (AI)^89^, was used to automate the process. Due to the limited size of our manually-curated training dataset (n = 790), we utilized MobileNetV2, a lightweight convolutional neural network (CNN) architecture that has been widely adopted for image classification tasks, as it repurposes knowledge gained from training on large, general datasets to improve performance on smaller, domain-specific datasets^90^. The pipeline, which we call Img-Cat, categorized CCM images into seven distinct classes based on growth, contamination status, and phenotypic complexity (Figure S3). The trained model had a validation accuracy of 93.22% and was saved for use in class prediction for consistent performance. When applied to the entire data set of 12,465 images, colonies were sorted by the pipeline and manually verified for accurate classification.

#### PSH

PSH was quantified in a two-step process. First, Img-Cat (described above) was trained with 758 manually-curated images to categorize test images into four classes: colony with PSH, colony with no PSH, contamination, and no growth (Figure S4). In total, 3744 images were categorized through the CNN pipeline, which had an accuracy above 90%, then manually verified for correct categorization. Next, the images with PSH were analyzed using a modified version of a previously published Python pipeline^50^, which we call Eclipse 2.0. Images were converted to grayscale, and a background threshold was determined using a smoothed histogram to exclude non-colony regions. Pixels with brightness exceeding a defined “white threshold” were marked as colony pixels, while intermediate-intensity pixels were classified separately as pseudohyphae pixels. A binary mask was generated to isolate colony regions, followed by connected component analysis and morphological operations to refine the colony boundary. The final processed image was analyzed by calculating the ratio of pseudohyphae pixels to core colony pixels, expressed as the PSH percentage (Figure S5).

#### Invasive Growth

A custom Python pipeline was used to process pre-washed and post-washed images of the SLAD 2X plates by applying a brightness threshold to the images to divide the pixels in an image into two clusters: (1) colony, (2) background. IG was quantified by dividing the number of colony pixels in the post-wash processed image by the number of colony pixels in the pre-wash processed image (Figure S6).

#### Plastic Adherence

A custom Python pipeline was used to process the images of the bottom of PA wells after washing away culture and non-adherent cells. A threshold was applied to the image to divide the image binarily: (1) yeast cells, (2) background. PA was quantified using the number of adhering yeast cells (white pixels) divided by the total number of pixels in the well, represented by the circles in Figure S7.

### Phenotypic Data Analysis

All phenotypic data were processed and analyzed in R^91^, and unless otherwise noted, figures were generated using ggplot2^92^. The control plates were compared among the three assays to verify that there were no systematic differences among the assays/ batches of plates (Figure SF). First, each phenotype was analyzed using an ordinary least squares regression (OLS) using the *lm()* function. The average of the three replicates was modeled with either the ecotype or clade as the predictor. Next, to test for the effect of shared evolutionary history, phylogenetic signal was calculated using Pagel’s Lambda (λ). The phylogenetic tree of the *S. cerevisiae* strains^18^ was made dichotomous using *multi2di()* in the *ape* package^93^ and the input matrix containing multicellular trait values was aligned to it. Any missing values were excluded prior to analysis; data on CCM in YPD medium were excluded from correlational analyses, as YPD serves as a nutrient-rich control condition. λ was calculated using *phylosig* from the *phytools* package^94^. To measure the two-sided correlation between traits while accounting for the influence of shared ancestry, trait values were transformed into phylogenetically independent contrasts (PIC) using Felsenstein’s method under a Brownian motion model^95^ with the *pic()* function from the *ape* package. Pairwise correlations were calculated using *cor()* and visualized using *corrplot*^96^. Finally, principal component analyses (PCA) were performed on phenotype averages and on PIC data using *prcomp()* to identify the major axes of variation across strains. The phylogenetic tree was visualized and annotated using iTOL (v7)^97^.

### Genome Wide Associate Studies (GWAS)

GWAS was performed to identify SNPs associated with multicellular phenotypes. The filtered genotype files were downloaded^18^ and converted to binary format using PLINK (v.1.90)^98^. Linear mixed-model association analysis was performed using FaST-LMM (v.0.6.10)^99^. The model expects phenotypes to have Gaussian residuals, thus the ordinal, categorical phenotypes were subject to a Yeo-Johnson transformation^100,101^ and the continuous, numerical phenotypes to a quantile transformation^100^. The significant p-value thresholds for individual phenotypes were estimated by permutation tests using the lowest 5th percentile of the p-values calculated over 100 randomizations. To correct for population structure, linkage disequilibrium (LD) pruning and principal component analysis were performed using PLINK. The cumulative variance explained showed small increments after the sixth principal component; therefore, the first 6 PCs were included in the covariate matrix used in the association and permutation tests. The genomic control inflation factor (λ_GC_) was computed using FaST-LMM. SNPs with significant p-values were mapped to coding genes using a custom Python pipeline and the annotated *Saccharomyces cerevisiae* genome (SGD R64-4-1)^70^. This pipeline was verified by, and further variant information was obtained, using snpEff 5.2c^102^. The 1011Matrix.gvcf was filtered by SNP chromosomes and positions using bcftools v.1.21^103^. Narrow sense heritability was estimated with variants on chromosomes 1-16 and phenotypic variance under the null model by FastLMM. The overlap in genes was determined using the VennDiagram package^104^ and visualized using ggVennDiagram^105^.

### Gene Ontology (GO) Enrichment Analysis

To gain insight into the biological functions and pathways enriched among genes associated with multicellular trait formation, GO enrichment analysis was conducted using ShinyGO 0.8^106^. The input gene list for GO analysis was compiled using the non-repetitive genes mapped by SNPs identified through GWAS. The parameters used for the analyses were: FDR cutoff: 0.05; Use pathway DB for gene counts; # of pathways to show: 20; Min pathway size: 2; Max pathway size: 5000.

### Virulence Assays

Strains for this experiment were selected for their ability to be: (1) multicellular in all traits, (2) highly multicellular in a single trait/suite of correlated traits, or (3) non-multicellular in all traits. Strains identified for each category were then re-assayed individually to verify their phenotypes and a final panel of 19 strains was generated (Table S5). Overnight yeast cultures grown in 10 ml YPD were washed twice and resuspended in sterile diH_2_O to a concentration of 1 x 10^9^ cells/ml using hemocytometer counts. Flocculated cultures (BBD and ALQ) were suspended in 1 ml of deflocculation buffer before washing. Four microliters of yeast or sterile diH_2_O were injected into the final posterior proleg of *Galleria mellonella* larvae using a Hamilton PB600-1 Repeating Dispenser with a 28-gauge 2-inch needle (Hamilton 7803-01). To control for potential variation, each of three *G. mellonella* shipments (Vanderhosrt Wholesale Inc.) was challenged with all yeast strains over a 2-day period. Larvae with a healthy appearance and weighing 180 mg (± 20 mg) were dispersed in groups of 20 into petri dishes. Each strain was injected into two plates (40 larvae); each day, one control plate was injected with sterile diH_2_O at the beginning and end of the assay. Between each 20-larvae group/petri dish, the injector was sterilized by twice drawing up and discarding ∼90 µl of 70% ethanol, then five times ∼100 µl of sterile diH_2_O to ensure no ethanol remained in the dispenser. Each yeast strain was injected into a total of 120 larvae (40 larvae/strain/shipment). Following injections, petri dishes containing larvae and sawdust from shipment containers were incubated at 30C in a ventilated box and monitored daily for 7 days. Larvae that were melanized and no longer responded to tactile stimulation were considered dead and removed from the plate, as were larvae beginning to pupate. Data were analyzed using the *survival*^107^ and *coxme*^108^ packages in R. All larvae started at t = 0; death was recorded on the day larvae were removed from the population, and pupated larvae were censored as they were progressing to the next life stage. The Cox Mixed Effects Model was used to analyze the data with strain and assay as random effects, and multicellular status status as a fixed effect.

## Supporting information

Supplemental Methods and Data Analysis

## DATA AVAILABILITY

Phenotypic data and images are available at https://y1000multicellularitymurphy.wm.edu/. Image analysis and GWAS code are available at https://github.com/gdzhong03/1011_multicellularity/tree/main.

## ACKNOWLEDGEMENTS

We thank Paul Rowley and Gianni Liti for access to the 1011 genomes collection. This research was supported by grants NIH-R15GM152938 and NSF DEB-2151034 to HAM, William & Mary Charles Center fellowships to DZ, LIE, AS, MVC, and AA.

## Supplementary Figure Legends and Tables

**Figure S1:**
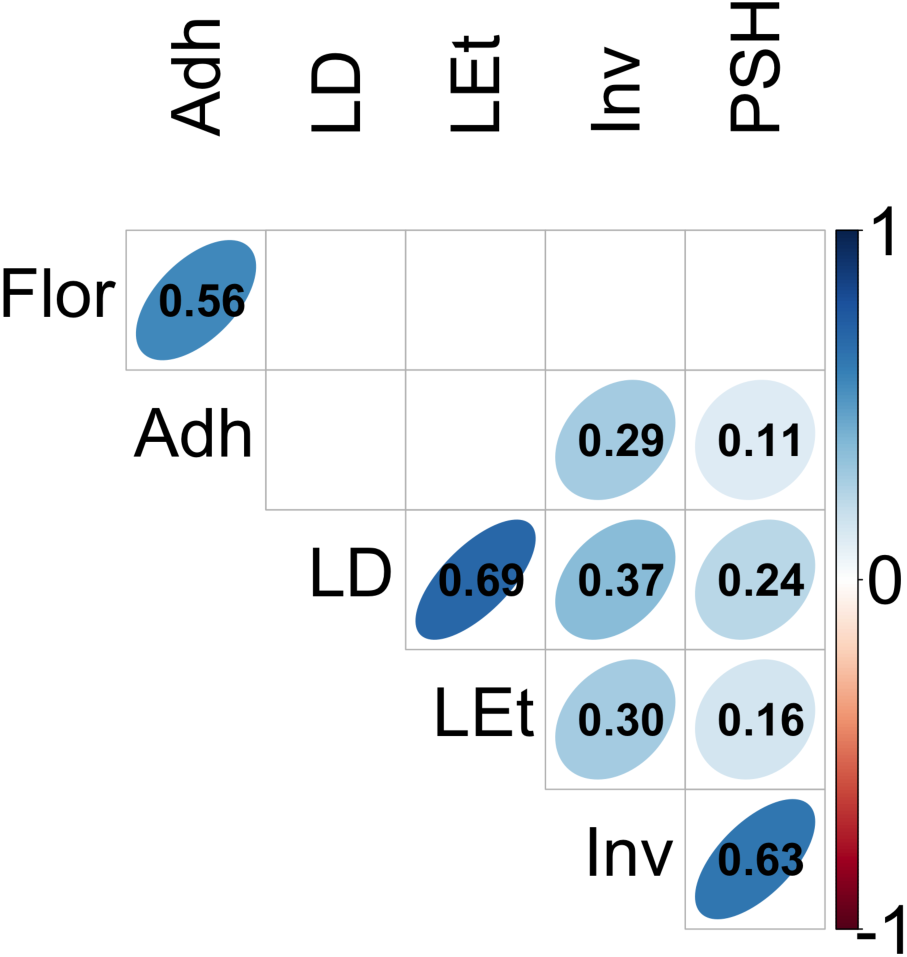
Pearson correlations between phenotypes without first correcting for phylogenetic relationships.

**Figure S2:**
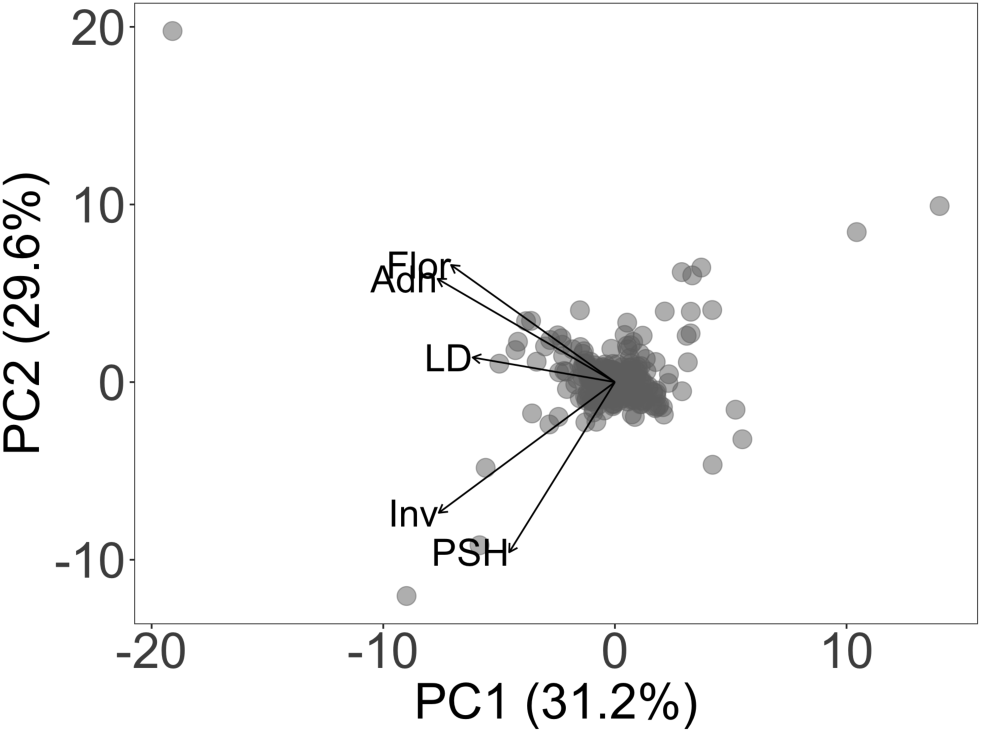
A principal component analysis on the pic-corrected phenotypes. PC1 (31.2%): -0.472*Flor - 0.409*LD - 0.507*Inv - 0.304*PSH - 0.510*Adh. PC2 (29.6%): -0.440*Flor - 0.093*LD + 0.490*Inv + 0.638*PSH - 0.387*Adh.

**Figure S3:**
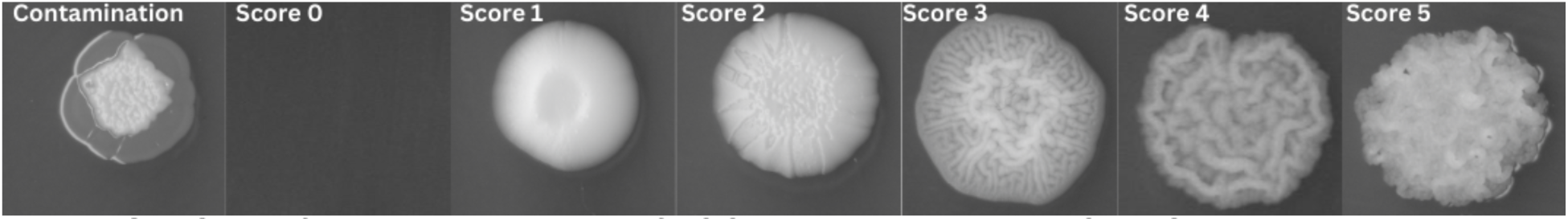
Quantification scale to classify CCM images using ImgCat . Score 0: no growth, score 1: colony with no CCM, score 2: colony with low levels of CCM, score 3: colony with moderate levels of CCM, score 4: colony with high levels of CCM, score 5: colony with extreme levels of CCM.

**Figure S4:**
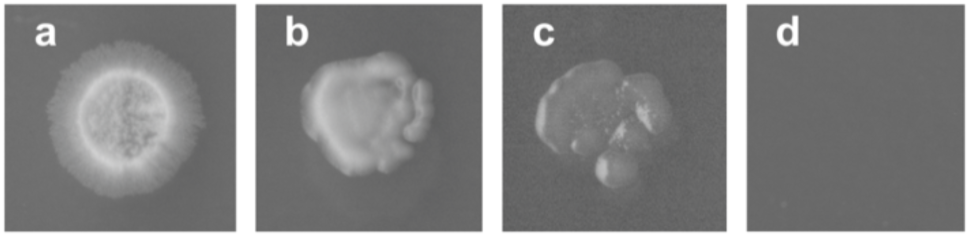
Classification categories for PSH. (a) PSH, (b) no PSH, (c) contamination, (d) no growth.

**Figure S5:**
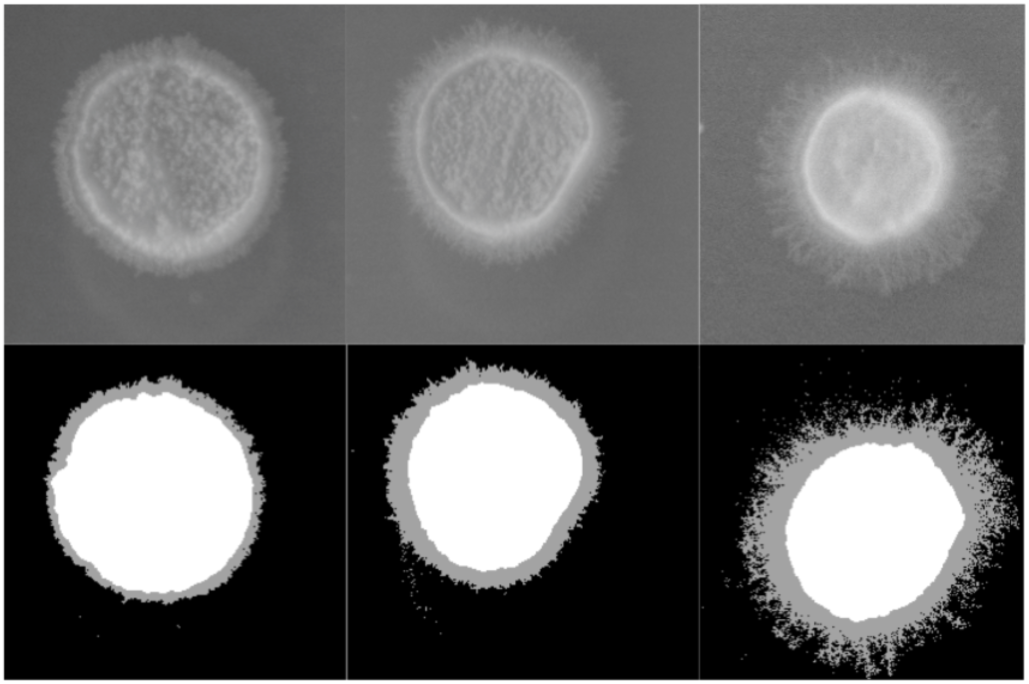
Illustration of Eclipse 2.0. PSH % is calculated using the number of pixels in pseudohyphae segments (gray) divided by the number of pixels in the main colony segment (white). PSH % from left to right: 24.76 %, 48.25 %, 95.10 %.

**Figure S6:**
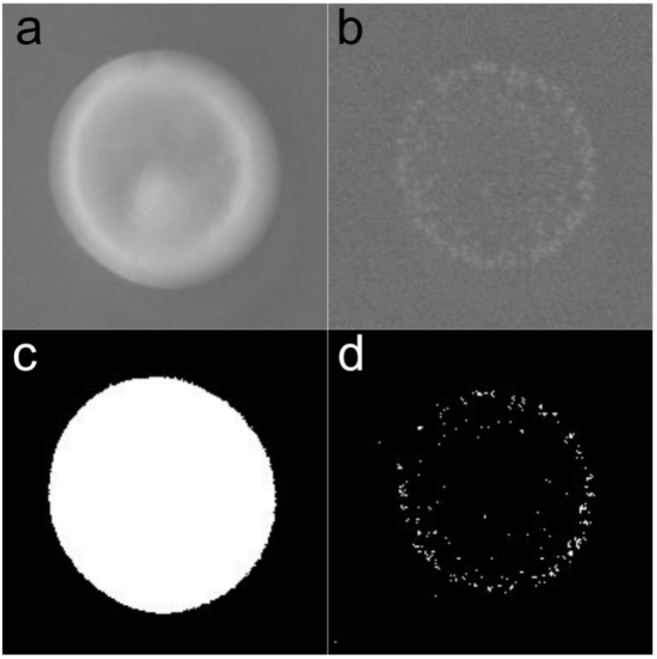
Invasive growth quantification: (a) Preprocessed, unwashed colony; (b) preprocessed, washed colony; (c) processed, unwashed colony; (d) processed, washed colony.

**Figure S7:**
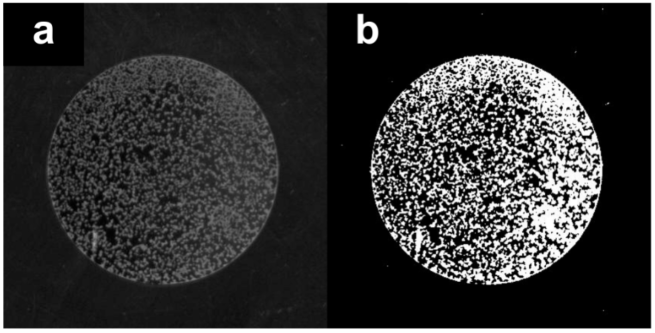
PA quantification: (a) Preprocessed image of one well of a washed 96-well plate containing adherent *S. cerevisiae* cells; (b) processed by computational pipeline.

**Table S1:** Number and percentage of isolates exhibiting the phenotypes in each ecotype isolation category. The values underneath the phenotype names list the cut-off values for an isolate to be considered expressing the phenotype.

| Ecotype | N | YPD<br>(≥1.67) | D<br>(≥1.67) | LEt<br>(≥1.67) | Flor<br>(≥0.67) | PSH<br>(>0) | Adh<br>(>2) | Invasive<br>(>1) |
| --- | --- | --- | --- | --- | --- | --- | --- | --- |
| Human | 135 | 29<br>(21.5%) | 64<br>(47.5%) | 73<br>(54%) | 10<br>(7.5%) | 68<br>(50%) | 44<br>(33%) | 52<br>(38.5%) |
| Domesticated | 579 | 75<br>(13%) | 140<br>(24%) | 162<br>(30%) | 148<br>(25.5%) | 218<br>(37.5%) | 215<br>(37%) | 167<br>(29%) |
| Wild | 245 | 34<br>(14%) | 68<br>(28%) | 75<br>(30.5%) | 31<br>(12.5%) | 97<br>(39.5%) | 76<br>(31%) | 71<br>(29%) |
| Unknown | 39 | 9<br>(23%) | 10<br>(25.5%) | 13<br>(33.5%) | 6<br>(15.5%) | 15<br>(38.5%) | 10<br>(26%) | 11<br>(28%) |
| Total | 998 | 147<br>(14.5%) | 282<br>(28%) | 328<br>(32%) | 195<br>(19.5%) | 398<br>(40%) | 345<br>(34.5%) | 301<br>(30%) |

**Table S2:** Relationship between isolation ecotype and the multicellular phenotypes. Data were analyzed using an ordinary least-squares regression with the “wild” ecotype as the reference. Each cell contains the estimate of the coefficient with the standard error in parentheses, and the *p*-value below.

|  | CCM-LD | CCM-LEt | CCM-YPD | Flor | PA | PSH | IG |
| --- | --- | --- | --- | --- | --- | --- | --- |
| adj. R<sup>2</sup> | **0.029**<br>**5.2×10<sup>-7</sup>** | **0.031**<br>**4.2×10<sup>-7</sup>** | 0.003<br>0.134 | **0.021**<br>**2.7×10<sup>-5</sup>** | **0.007**<br>**0.019** | 0.0004<br>0.3306 | **0.01**<br>**0.008** |
| Intercept | 1.39 (0.04)<br><2×10<sup>-16</sup> | 1.44 (0.05)<br><2×10<sup>-16</sup> |  | 0.28 (0.06)<br>8.3×10<sup>-7</sup> | 3.32 (0.6)<br>3.8×10<sup>-8</sup> |  | 8.44 (1.39)<br>1.9×10<sup>-9</sup> |
| domestic | -0.09 (0.047)<br>0.0552 | -0.08 (0.055)<br>0.151 |  | **0.22 (0.07)**<br>**0.0016** | **1.8 (0.72)**<br>**0.012** |  | 0.52 (1.66)<br>0.753 |
| clinical | **0.25 (0.067)**<br>**0.0002** | **0.32 (0.078)**<br>**4.1×10<sup>-5</sup>** |  | -0.15 (0.10)<br>0.12 | **2.14 (1.01)**<br>**0.034** |  | **7.25 (2.34)**<br>**0.0019** |
| UNK | -0.05 (0.11)<br>0.6325 | -0.01 (0.12)<br>0.934 |  | -0.04 (0.15)<br>0.82 | -1.12 (1.62)<br>0.488 |  | -0.48 (3.93)<br>0.902 |

**Table S3:** Relationship between clade/subclade and the multicellular phenotypes. Clades are listed with the number of isolates in the clade. Data were analyzed using an ordinary least-squares regression with the 0.8 Mixed Origin clade (n = 72) as the reference. Each cell contains the estimate of the coefficient with the standard error in parentheses, and the *p*-value below. Bold and shading indicate significance.

|  | CCM-LD | CCM-LEt | CCM-YPD | Flor | PA | PSH | IG |
| --- | --- | --- | --- | --- | --- | --- | --- |
| adj. R<sup>2</sup> | **0.1555**<br>**<2×10<sup>-16</sup>** | **0.1406**<br>**<2×10<sup>-16</sup>** | **0.0346**<br>**0.0004664** | **0.2309**<br>**<2×10<sup>-16</sup>** | **0.258**<br>**<2×10<sup>-16</sup>** | **0.1491**<br>**<2×10<sup>-16</sup>** | **0.2437**<br>**2×10<sup>-16</sup>** |
| Intercept | 1.38 (0.07)<br><2×10<sup>-16</sup> | 1.53 (0.08)<br><2×10<sup>-16</sup> | 1.17 (0.05)<br><2×10<sup>-16</sup> | 0.23 (0.09)<br>0.01324 | 1.03 (0.96)<br>0.2811 | 10.62 (2.2)<br>9.8×10<sup>-7</sup> | 4.97 (2.27)<br>0.029 |
| **01 Wine Euro WE (267)** | **-0.19 (0.08)**<br>**0.016** | **-0.26 (0.09)**<br>**0.006** | -0.08 (0.06)<br>0.13 | 0.18 (0.11)<br>0.09 | **2.54 (1.08)**<br>**0.019** | 4.28 (2.42)<br>0.077 | **5.7 (2.55)**<br>**0.26** |
| **01.W1 WE - Wild (18)** | -0.10 (0.15)<br>0.52 | -0.03 (0.18)<br>0.86 | -0.06 (0.11)<br>0.56 | **2.14 (0.21)**<br>**<2×10<sup>-16</sup>** | **11.33(2.14)**<br>**1.4×10<sup>-7</sup>** | -5.42 (4.65)<br>0.24 | -1.85 (4.9)<br>0.71 |
| **01.W2 WE- Clinic. (13)** | -0.33 (0.17)<br>0.06 | **-0.42 (0.21)**<br>**0.047** | -0.17 (0.12)<br>0.16 | -0.23 (0.24)<br>0.33 | -0.29 (2.44)<br>0.91 | -7.46 (5.31)<br>0.16 | -3.08 (5.6)<br>0.58 |
| **01.W3 WE- Clinic./ S. boulardi(23)** | **0.41 (0.14)**<br>**0.0036** | 0.25 (0.16)<br>0.12 | 0.03 (0.10)<br>0.77 | -0.23 (0.19)<br>0.22 | 0.32 (1.94)<br>0.87 | **-10.12 (4.31)**<br>**0.019** | -4.93 (4.54)<br>0.28 |
| **01.W4 WE- Georg. (39)** | -0.06 (0.12)<br>0.60 | -0.26 (0.14)<br>0.071 | -0.05 (0.08)<br>0.53 | **1.56 (0.16)**<br>**<2×10<sup>-16</sup>** | **14.38(1.61)**<br>**<2×10<sup>-16</sup>** | **-8.74 (3.63)**<br>**0.016** | 4.75 (3.82)<br>0.21 |
| **02 Alpechin (17)** | -0.21 (0.16)<br>0.185 | -0.23 (0.19)<br>0.234 | 0.04 (0.11)<br>0.708 | -0.23 (0.24)<br>0.326 | 0.01 (2.29)<br>0.997 | -6.75 (4.84)<br>0.164 | -4.59 (5.06)<br>0.365 |
| **03 Brazil Bioethanol (35)** | -0.15 (0.12)<br>0.243 | **-0.28 (0.15)**<br>**0.055** | 0.16 (0.09)<br>0.0673 | -0.07 (0.18)<br>0.699 | 1.17 (1.75)<br>0.505 | **-9.74 (4.05)**<br>**0.016** | -4.80 (4.31)<br>0.265 |
| **04 Oak Mediterr (8)** | -0.25 (0.22)<br>0.249 | -0.28 (0.25)<br>0.268 | 0.08 (0.15)<br>0.621 | -0.23 (0.33)<br>0.4772 | 0.48 (3.16)<br>0.878 | -4.42 (6.66)<br>0.507 | -0.46 (6.97)<br>0.948 |
| **05 French Dairy (32)** | 0.02 (0.13)<br>0.879 | -0.21 (0.15)<br>0.155 | -0.03 (0.09)<br>0.750 | **0.48 (0.19)**<br>**0.0103** | **24.63 (1.8)**<br>**<2×10<sup>-16</sup>** | **20.25 (3.77)**<br>**9.8×10<sup>-8</sup>** | **48.14 (3.98)**<br>**<2×10<sup>-16</sup>** |
| **06 African beer (20)** | 0.24 (0.15)<br>0.118 | **-0.39 (0.18)**<br>**0.027** | -0.06 (0.10)<br>0.584 | **0.99 (0.22)**<br>**9.1x10<sup>-6</sup>** | **5.9 (2.15)**<br>**0.004** | **17.83 (4.56)**<br>**9.8x10<sup>-5</sup>** | **16.42 (4.81)**<br>**0.000664** |
| **07 Mosaic beer (21)** | -0.13 (0.15)<br>0.386 | 0.024 (0.18)<br>0.893 | -0.03 (0.11)<br>0.753 | 0.23 (0.22)<br>0.291 | **5.2 (2.01)**<br>**0.009** | -2.17 (4.54)<br>0.633 | 5.88 (4.75)<br>0.216 |
| **09 Mexican Agave (7)** | -0.33 (0.23)<br>0.156 | -0.49 (0.27)<br>0.073 | -0.13 (0.16)<br>0.439 | -0.23 (0.35)<br>0.503 | -0.68 (3.36)<br>0.840 | -7.88 (7.08)<br>0.266 | -2.72 (7.40)<br>0.713 |
| **10 Human Fr. Guiana (31)** | **1.11 (0.126)**<br>**<2x10<sup>-16</sup>** | **1.02 (0.15)**<br>**1.38x10<sup>-11</sup>** | **0.36 (0.09)**<br>**6.94x10<sup>-5</sup>** | -0.19 (0.19)<br>0.315 | 2.64 (1.82)<br>0.148 | **22.93 (4.0)**<br>**9.8x10<sup>-8</sup>** | **32.86 (4.22)**<br>**1.8x10<sup>-14</sup>** |
| **11 Ale beer (17)** | -0.22 (0.16)<br>0.165 | **-0.37 (0.18)**<br>**0.043** | 0.08 (0.11)<br>0.467 | 0.05 (0.23)<br>0.841 | 1.10 (2.24)<br>0.623 | 1.83 (5.24)<br>0.726 | 6.01 (5.48)<br>0.272 |
| **12 W. Africa cocoa (13)** | **-0.38 (0.18)**<br>**0.0395** | **-0.50 (0.21)**<br>**0.0181** | -0.17 (0.13)<br>0.177 | -0.07 (0.27)<br>0.812 | -0.7 (2.65)<br>0.792 | 5.94 (5.59)<br>0.288 | -4.38 (5.84)<br>0.454 |
| **13 African palm wine (28)** | -0.04 (0.13)<br>0.74 | -0.013 (0.16)<br>0.935 | -0.06 (0.09)<br>0.552 | -0.09 (0.20)<br>0.649 | -0.23 (1.89)<br>0.904 | 5.51(4.07)<br>0.176 | 0.35 (4.25)<br>0.935 |
| **14 CHNIII (2)** | 0.79 (0.42)<br>0.0598 | 0.47 (0.49)<br>0.335 | -0.17 (0.30)<br>0.556 | -0.23 (0.63)<br>0.712 | 1.7 (6.08)<br>0.780 | -10.62 (17.9)<br>0.554 | -4.95 (18.75)<br>0.792 |
| **15 CHNII (2)** | -0.38 (0.42)<br>0.369 | -0.53 (0.49)<br>0.274 | -0.17 (0.30)<br>0.556 | -0.23 (0.63)<br>0.712 | -0.10 (6.08)<br>0.987 | -4.67 (12.78)<br>0.715 | -4.58 (13.35)<br>0.732 |
| **16 CHNI (1)** | 0.29 (0.59)<br>0.623 | -0.53 (0.68)<br>0.436 | -0.17 (0.41)<br>0.675 | -0.23 (0.88)<br>0.792 | -0.50 (8.55)<br>0.954 | -10.62 (17.9)<br>0.554 | -4.07 (18.75)<br>0.828 |
| **17 Taiwanese (3)** | -0.043 (0.35)<br>0.8996 | -0.198 (0.40)<br>0.620 | 0.16 (0.24)<br>0.511 | -0.23 (0.52)<br>0.653 | 0.31 (5.00)<br>0.951 | -10.62 (10.5)<br>0.313 | -4.95 (10.98)<br>0.653 |
| **18 Far East Asia (9)** | -0.23 (0.21)<br>0.2698 | -0.45 (0.25)<br>0.0776 | -0.10 (0.15)<br>0.494 | 0.10 (0.31)<br>0.742 | 0.69 (3.00)<br>0.818 | 0.93 (6.66)<br>0.889 | -2.82 (6.97)<br>0.686 |
| **19 Malaysian (6)** | -0.38 (0.35)<br>0.275 | -0.53 (0.40)<br>0.184 | -0.17 (0.24)<br>0.474 | -0.23 (0.40)<br>0.567 | 0.54 (3.93)<br>0.890 | -10.62 (10.5)<br>0.313 | -1.72 (10.98)<br>0.876 |
| **20 CHNV (2)** | -0.38 (0.42)<br>0.369 | -0.53 (0.49)<br>0.274 | -0.17 (0.30)<br>0.556 | -0.23 (0.63)<br>0.712 | 1.65 (6.08)<br>0.786 | -10.62 (12.8)<br>0.406 | -4.74 (13.35)<br>0.723 |
| **21 Ecuador (10)** | -0.38 (0.198)<br>0.057 | -0.50 (0.23)<br>0.0306 | -0.17 (0.14)<br>0.212 | -0.23 (0.30)<br>0.432 | 0.06 (2.86)<br>0.983 | -10.62 (6.22)<br>0.088 | -4.9 (6.61)<br>0.459 |
| **22 Russian (4)** | -0.38 (0.30)<br>0.210 | -0.53 (0.35)<br>0.128 | -0.17 (0.21)<br>0.411 | -0.23 (0.45)<br>0.606 | 0.93 (4.36)<br>0.832 | -10.62 (9.17)<br>0.247 | -4.93 (9.58)<br>0.607 |
| **23 Oak N America (13)** | -0.22 (0.18)<br>0.207 | -0.25 (0.21)<br>0.226 | 0.01 (0.12)<br>0.964 | -0.18 (0.26)<br>0.494 | 0.34 (2.56)<br>0.895 | -8.6 (5.4)<br>0.112 | -4.58 (5.65)<br>0.417 |
| **24 Asian islands (11)** | 0.26 (0.19)<br>0.172 | -0.07 (0.23)<br>0.779 | -0.08 (0.13)<br>0.534 | 0.01 (0.28)<br>0.969 | **7.23 (2.66)**<br>**0.006** | **11.66 (5.70)**<br>**0.041** | **15.83 (6.01)**<br>**0.009** |
| **25 Sake (47)** | 0.20 (0.11)<br>0.079 | **0.51 (0.13)**<br>**9.55x10<sup>-6</sup>** | 0.15 (0.08)<br>0.064 | **0.37 (0.15)**<br>**0.013** | **4.69 (1.52)**<br>**0.002** | -6.02 (3.4)<br>0.075 | -1.78 (3.60)<br>0.621 |
| **26 Asian ferment (39)** | 0.057 (0.12)<br>0.627 | 0.14 (0.14)<br>0.3324 | 0.12 (0.08)<br>0.165 | -0.17 (0.18)<br>0.332 | 1.6 (1.7)<br>0.345 | **-9.45 (3.66)**<br>**0.0099** | -4.31 (3.89)<br>0.269 |
| **99 Other (46)** | 0.09 (0.11)<br>0.43 | -0.04 (0.13)<br>0.75 | 0.06 (0.08)<br>0.46 | -0.17 (0.15)<br>0.25 | 0.52 (1.53)<br>0.73 | -0.94 (3.56)<br>0.79 | -2.57 (3.76)<br>0.49 |
| **M1 Mosaic Reg 1 (17)** | -0.22 (0.16)<br>0.16 | -0.32 (0.18)<br>0.09 | -0.12 (0.11)<br>0.30 | -0.19 (0.21)<br>0.37 | 0.56 (2.19)<br>0.80 | 2.06 (4.88)<br>0.67 | -0.31 (5.14)<br>0.95 |
| **M2 Mosaic Reg 2 (19)** | 0.13 (0.16)<br>0.40 | -0.18 (0.18)<br>0.33 | -0.12 (0.11)<br>0.30 | -0.21 (0.20)<br>0.29 | 0.14 (2.09)<br>0.95 | 9.22 (5.0)<br>0.066 | 9.43 (5.28)<br>0.074 |
| **M3 Mosaic Reg 3 (108)** | 0.14 (0.09)<br>0.12 | -0.01 (0.11)<br>0.92 | 0.07 (0.06)<br>0.29 | 0.06 (0.12)<br>0.61 | 5.23 (1.23)<br>**2.4x10<sup>-5</sup>** | 2.51 (2.76)<br>0.36 | **8.3 (2.92)**<br>**0.0045** |

**Table SX:** Relationship between the ecological origin of the clade and the multicellular phenotypes. Data were analyzed using an ordinary least-squares regression with the Nature category as the reference. Each cell contains the estimate of the coefficient with the standard error in parentheses, and the *p*-value below.

|  | CCM-LD | CCM-LEt | CCM-YPD | Flor | PA | PSH | IG |
| --- | --- | --- | --- | --- | --- | --- | --- |
| adj. R<sup>2</sup> | **0.134**<br>**<2x10<sup>-16</sup>** | **0.1579**<br>**<2x10<sup>-16</sup>** | **0.04269**<br>**4.4x10<sup>-6</sup>** | **0.06613**<br>**3.05x10<sup>-10</sup>** | **0.1148**<br>**<2x10<sup>-16</sup>** | **0.0927**<br>**5.1x10<sup>-14</sup>** | **0.2176**<br>**2x10<sup>-16</sup>** |
| Intercept | 1.48 (0.08)<br>**<2x10<sup>-16</sup>** | 1.37 (0.10)<br>**<2x10<sup>-16</sup>** | 1.25 (0.06)<br>**<2x10<sup>-16</sup>** | 0.18 (0.12)<br>0.134 | 3.85 (1.24)<br>0.002 | 12.88 (2.6)<br>1.2x10<sup>-5</sup> | 10.2 (2.74)<br>0.0002 |
| Bakery (34) | 0.04 (0.13)<br>0.778 | 0.27 (0.15)<br>0.080 | 0.04 (0.09)<br>0.640 | -0.02 (0.19)<br>0.933 | -1.78 (1.96)<br>0.364 | -2.09 (4.3)<br>0.625 | 1.29 (4.43)<br>0.771 |
| Beer (54) | -0.13 (0.11)<br>0.272 | -0.12 (0.13)<br>0.378 | -0.09 (0.08)<br>0.288 | **0.52 (0.17)**<br>**0.0018** | -0.01 (1.69)<br>0.996 | 6.44 (3.62)<br>0.076 | 0.20 (3.76)<br>0.958 |
| Bioethanol (27) | -0.24 (0.14)<br>0.089 | -0.11 (0.16)<br>0.511 | 0.19 (0.1)<br>0.056 | -0.08 (0.21)<br>0.685 | -2.77 (2.11)<br>0.190 | -12.7 (5.0)<br>0.012 | -9.99 (5.20)<br>0.055 |
| Cider (17) | **-0.34 (0.17)**<br>**0.044** | -0.08 (0.19)<br>0.667 | -0.19 (0.12)<br>0.114 | -0.16 (0.24)<br>0.504 | -2.10 (2.48)<br>0.397 | -2.57 (5.5)<br>0.640 | -9.60 (5.72)<br>0.094 |
| Dairy (27) | -0.10 (0.14)<br>0.469 | -0.05 (0.16)<br>0.739 | -0.08 (0.10)<br>0.439 | **0.52 (0.21)**<br>**0.0123** | **19.0 (2.11)**<br>**<2x10<sup>-16</sup>** | **17.4 (4.36)**<br>**7.1x10<sup>-5</sup>** | **46.03(4.53)**<br>**<2x10<sup>-16</sup>** |
| Distillery (29) | -0.12 (0.14)<br>0.372 | 0.08 (0.16)<br>0.594 | -0.01 (0.10)<br>0.942 | -0.10 (0.20)<br>0.613 | -2.75 (2.06)<br>0.183 | -4.0 (4.27)<br>0.349 | 0.19 (4.43)<br>0.966 |
| Ferment (36) | -0.21 (0.13)<br>0.107 | -0.05 (0.15)<br>0.756 | -0.14 (0.09)<br>0.124 | -0.07 (0.19)<br>0.705 | -0.11 (1.93)<br>0.956 | -2.36(4.04)<br>0.559 | -5.45 (4.19)<br>0.194 |
| Flower (14) | -0.27 (0.18)<br>0.128 | 0.10 (0.20)<br>0.613 | -0.18 (0.12)<br>0.153 | **0.84 (0.26)**<br>**0.0014** | 2.51 (2.67)<br>0.348 | **-12.8 (5.5)**<br>**0.0195** | -10.04(5.72)<br>0.079 |
| Fruit (45) | 0.09 (0.12)<br>0.432 | 0.26 (0.14)<br>0.056 | -0.02 (0.08)<br>0.823 | 0.05 (0.18)<br>0.794 | -1.60 (1.81)<br>0.376 | 3.39 (3.81)<br>0.374 | 3.37 (3.96)<br>0.395 |
| Human Feces (31) | **0.87 (0.13)**<br>**1x10<sup>-10</sup>** | **1.17 (0.15)**<br>**1x10<sup>-13</sup>** | **0.22 (0.09)**<br>**0.0208** | -0.04 (0.20)<br>0.828 | 1.38 (2.02)<br>0.495 | **16.6 (4.36)**<br>**0.000152** | **22.32 (4.53)**<br>**1x10<sup>-6</sup>** |
| Human Clinical (104) | -0.07 (0.1)<br>0.479 | 0.15 (0.12)<br>0.194 | -0.09 (0.07)<br>0.201 | -0.05 (0.15)<br>0.729 | 1.69 (1.52)<br>0.266 | -3.85 (3.2)<br>0.229 | 0.951 (3.32)<br>0.775 |
| Industrial (30) | -0.21 (0.14)<br>0.117 | -0.04 (0.16)<br>0.801 | -0.06 (0.10)<br>0.499 | 0.02 (0.20)<br>0.933 | -0.84 (2.04)<br>0.680 | -3.68(4.36)<br>0.399 | -6.80 (4.53)<br>0.134 |
| Insect (20) | -0.21 (0.16)<br>0.179 | -0.11 (0.18)<br>0.537 | -0.11 (0.11)<br>0.327 | 0.15 (0.23)<br>0.514 | -0.06 (2.34)<br>0.981 | -2.74(5.23)<br>0.601 | -2.08 (5.43)<br>0.702 |
| Lab (2) | **1.52 (0.42)**<br>**0.0003** | **1.96 (0.48)**<br>**4.9x10<sup>-5</sup>** | **0.75 (0.30)**<br>**0.0111** | -0.18 (0.63)<br>0.771 | -2.61 (6.39)<br>0.683 | -12.9(18.3)<br>0.481 | -10.2 (19.0)<br>0.592 |
| Palm wine (29) | -0.25 (0.14)<br>0.063 | -0.14 (0.16)<br>0.388 | -0.18 (0.10)<br>0.0601 | 0.04 (0.20)<br>0.862 | -2.42 (2.06)<br>0.241 | 1.39 (4.27)<br>0.746 | -0.38 (4.43)<br>0.932 |
| Sake (47) | 0.06 (0.12)<br>0.637 | **0.58 (0.14)**<br>**2.2x10<sup>-5</sup>** | 0.03 (0.08)<br>0.746 | **0.42 (0.18)**<br>**0.0175** | 1.89 (1.79)<br>0.291 | **-8.5 (3.77)**<br>**0.0243** | -7.34 (3.92)<br>0.0611 |
| Soil (38) | **-0.34 (0.13)**<br>**0.0063** | -0.10 (0.15)<br>0.513 | -0.15 (0.09)<br>0.084 | 0.001 (0.2)<br>0.995 | -1.65 (1.90)<br>0.386 | -5.16 (3.97)<br>0.195 | **-8.9 (4.13)**<br>**0.0311** |
| Tree (63) | **-0.26 (0.11)**<br>**0.0184** | -0.10 (0.13)<br>0.452 | -0.14 (0.08)<br>0.081 | 0.03 (0.16)<br>0.836 | -0.28 (1.67)<br>0.867 | -5.03 (3.52)<br>0.154 | -6.85 (3.66)<br>0.0614 |
| Unknown (28) | -0.26 (0.14)<br>0.0618 | -0.10 (0.16)<br>0.520 | -0.12 (0.10)<br>0.224 | 0.12 (0.21)<br>0.576 | -1.08 (2.08)<br>0.604 | 0.43 (4.42)<br>0.922 | -4.23 (4.59)<br>0.357 |
| Water (19) | **0.44 (0.16)**<br>**0.00597** | **0.87 (0.18)**<br>**2.7x10<sup>-7</sup>** | 0.09 (0.11)<br>0.450 | 0.29 (0.23)<br>0.215 | -1.36 (2.38)<br>0.569 | **18.1 (5.23)**<br>**0.00057** | **22.5 (5.43)**<br>**0.0000384** |
| Wine (248) | -0.27 (0.09)<br>0.0024 | -0.12 (0.11)<br>0.264 | **-0.16 (0.06)**<br>**0.0107** | **0.51 (0.13)**<br>**0.000143** | 1.92 (1.36)<br>0.159 | -1.94 (2.9)<br>0.503 | -3.75 (3.00)<br>0.213 |

**Table S4:** Pearson correlation between multicellular phenotypes, both with and without correction for phylogentically independent contrasts (PIC).

| Comparison | $r_{PIC}$ | $P\text{-value}_{PIC}$ | $r$ | $P\text{-value}$ |
| --- | --- | --- | --- | --- |
| Inv-PSH | **0.486** | **0** | **0.631** | **0** |
| Inv-LD | **0.156** | **$2.2 \times 10^{-6}$** | **0.373** | **0** |
| PSH-LD | 0.006 | 0.856 | **0.240** | **$1.8 \times 10^{-13}$** |
| Inv-LEt | **0.410** | **0** | **0.300** | **0** |
| PSH-LEt | **0.261** | **$8.8 \times 10^{-16}$** | **0.157** | **$1.8 \times 10^{-6}$** |
| LD-LEt | **0.518** | **0** | **0.693** | **0** |
| Inv-Flor | 0.065 | 0.0510 | 0.029 | 0.385 |
| PSH-Flor | **-0.111** | **$7.3 \times 10^{-4}$** | -0.009 | 0.789 |
| LD-Flor | **0.140** | **$2.1 \times 10^{-5}$** | 0.044 | 0.180 |
| LEt-Flor | **0.132** | **$6.0 \times 10^{-5}$** | **0.067** | **0.042** |
| Inv-Adh | **0.084** | **0.0115** | **0.292** | **0** |
| PSH-Adh | -0.050 | 0.129 | **0.112** | **0.001** |
| LD-Adh | **0.163** | **$7.4 \times 10^{-7}$** | **0.063** | **0.057** |
| LEt-Adh | **0.119** | **$3 \times 10^{-4}$** | **0.071** | **0.032** |
| Flor-Adh | **0.409** | **0** | **0.565** | **0** |

**Table S6.**
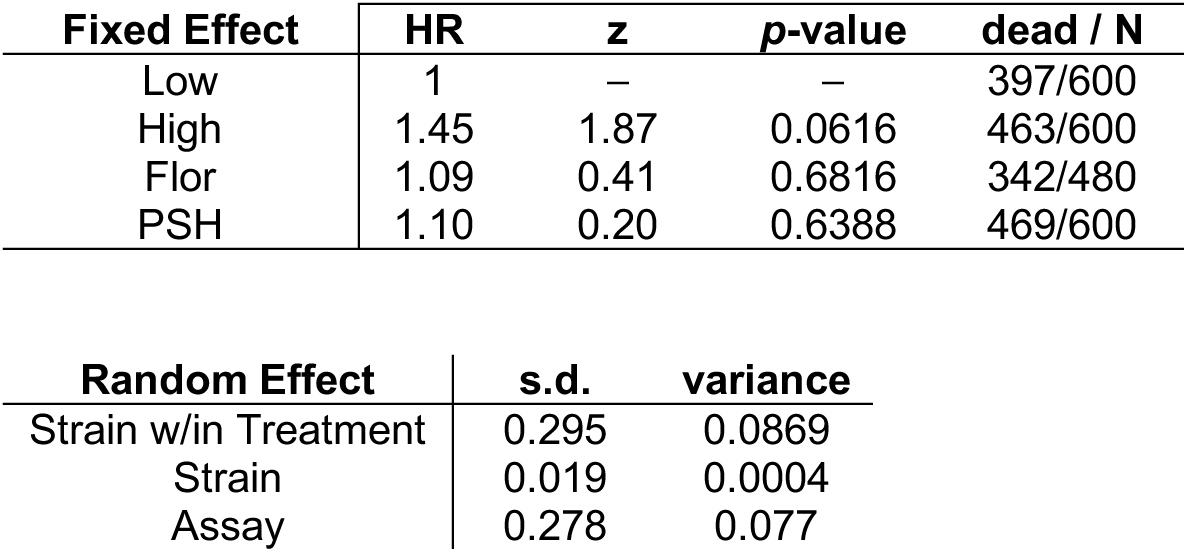
Results of a Cox Mixed Effect Model with multicellularity included as a fixed effect, and strains and assays included as random effects.

| Fixed Effect | HR | z | p-value | dead / N |
| --- | --- | --- | --- | --- |
| Low | 1 | — | — | 397/600 |
| High | 1.45 | 1.87 | 0.0616 | 463/600 |
| Flor | 1.09 | 0.41 | 0.6816 | 342/480 |
| PSH | 1.10 | 0.20 | 0.6388 | 469/600 |

| Random Effect | s.d. | variance |
| --- | --- | --- |
| Strain w/in Treatment | 0.295 | 0.0869 |
| Strain | 0.019 | 0.0004 |
| Assay | 0.278 | 0.077 |

**Table S7.** Results of a Cox Mixed Effect Model with strains as a fixed effect (to quantify differences among strains), and assays included as a random effect.

| Effect | HR | z | p-value | dead / N |
| --- | --- | --- | --- | --- |
| L1 | 1 | — | — | 91/120 |
| L2 | 1.083 | 0.54 | 0.5891 | 94/120 |
| L3 | **0.473** | **-4.48** | **7.54E-06** | **59/120** |
| L4 | **0.383** | **-5.52** | **3.39E-08** | **52/120** |
| L5 | **1.395** | **2.3** | **0.0214** | **101/120** |
| F1 | 1.003 | 0.02 | 0.9835 | 96/120 |
| F2 | 0.891 | -0.78 | 0.4355 | 91/120 |
| F3 | **0.695** | **-2.34** | **0.0193** | **76/120** |
| F4 | 0.858 | -0.99 | 0.3198 | 79/120 |
| P1 | 0.913 | -0.58 | 0.5611 | 82/120 |
| P2 | **0.497** | **-4.23** | **2.36E-05** | **61/120** |
| P3 | 1.274 | 1.69 | 0.0914 | 105/120 |
| P4 | 0.778 | -1.64 | 0.1006 | 81/120 |
| P5 | 1.022 | 0.15 | 0.8812 | 96/120 |
| H1 | 1.173 | 1.07 | 0.2829 | 106/120 |
| H2 | 1.186 | 1.16 | 0.2481 | 104/120 |
| H3 | 0.991 | -0.06 | 0.954 | 95/120 |
| H4 | 1.166 | 1.06 | 0.2884 | 100/120 |
| H5 | 1.202 | 1.27 | 0.2031 | 102/120 |

| Random Effect | s.d. | variance |
| --- | --- | --- |
| Assay | 0.278 | 0.077 |

### Separate Files

**Table S5:** List of strains used in the virulence assays.

**Table S8.** Complete list of significant SNPs found in all the GWASs.

**Table S9.** List of significant genes found in all the GWASs and the overlap between CCM-LD, Flor, PA, PSH, and IG.

**Table S10.** List of SNPs with overlap among phenotypes along with other SNPs in the same gene. Variants were queried for overlap among different combinations of the main 5 phenotypes: CCM-LD, Flor, PA, PSH, IG. When variants were identified, all other variants in the gene were listed for all the phenotypes (including LEt and YPD).

