## Supplemental Methods and Data Analysis for "Multicellularity and virulence in the budding yeast, *Saccharomyces cerevisiae*"

*Image Analysis*

CCM and PSH images needed to be categorized (0-5 for CCM, and presence/absence for PSH). We developed a pipeline to categorize the images, which we call Img-Cat.

To pre-process images, contrast was adjusted by a factor of 2, and to enhance the data variability, random brightness variations were introduced (max_delta=0.1). The images were normalized by rescaling the pixel intensity to a range of [0,1] and randomly split into training and validation datasets with an 8:2 ratio. The input image dimensions passed to MobileNetV2 are typically 224 x 224 x 3 (height, width, and color channels). After processing through the convolutional layers, the output is a 4 x 4 x 1280 feature map (i.e., a 4x4 grid of 1280 different feature maps, each capturing some learned pattern or feature from the image). Next, a GlobalAveragePooling2D layer was applied, which averages each of the 1280 feature maps across the 4x4 grid, converting them into 1280 1D values, thus reducing the spatial dimensions. while retaining the depth (the number of feature maps). These 1280 features were then connected to 512 neurons in a fully connected layer. The ReLU activation function was applied, ensuring all output values were positive, which introduced non-linearity and helped the model capture more complex patterns. To prevent overfitting, a Dropout layer was used, which randomly drops 50% of the neurons during training, forcing the model to generalize better by preventing it from relying too much on any one neuron. Finally, the remaining neurons were connected to the output layer, which contained 7 neurons (one for each class), and a softmax activation was applied. This converted the outputs into a probability distribution over the 7 classes, where the sum of all probabilities is 1. The model then classified the image into one of the 7 categories based on the highest probability. The training and validation accuracy were calculated separately and verified through manual examination.

### Data Analysis: *CCM- LEt and CCM-YPD*

As part of our phenotyping effort, we investigated complex colony morphology (CCM) on low ethanol, (YP-LEt, 0.05% ethanol) and rich medium (YPD, 2% glucose). The same analyses that are presented in the main text were performed for these media as well.

*Phenotype*
On low ethanol medium, 32% of the isolates exhibited CCM, which is similar to the low dextrose medium (28%, see the Main text). In contrast, on the rich YPD medium, only 14% of the isolates exhibited CCM (Figure S8A). This is not surprising, since carbon limitation is known to induce CCM. When CCM was analyzed by ecotype, on the LEt medium, clinical isolates had significantly greater CCM formation, while no ecotype was significant for YPD (Figure S8B, Table S2).

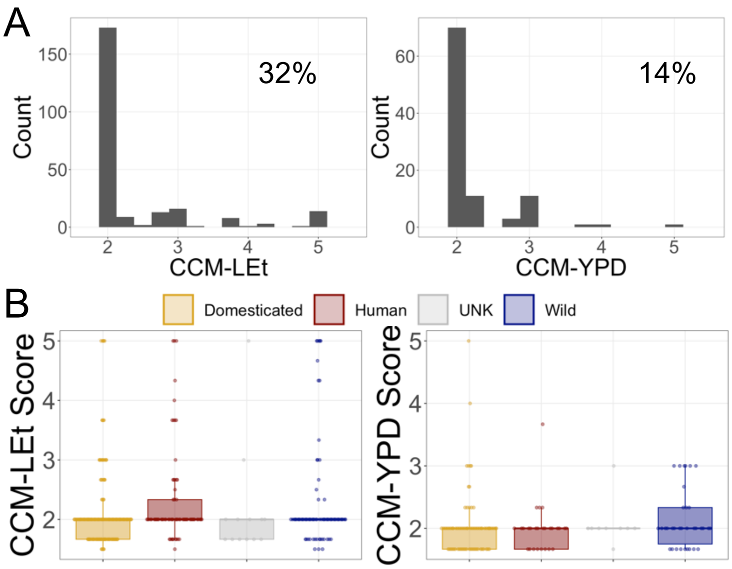


**Figure S8: Prevalence and distribution of CCM on LEt and YPD.**
**(A)** Phenotypic distributions for the subset of isolates that exhibit the phenotype (i.e., excluding the non-multicellular isolates). Strains had to score at least 1.67 to be included (i.e., at least two of the three replicates were scored at 2). The inset lists the percentage of isolates in the panel that exhibit the phenotype and on which the distribution is based. **(B)** Average phenotype values for all of the isolates in each ecotype category.

Next, the CCM data were analyzed by clade origin using an ordinary least squares regression (OLS). CCM on LEt showed significant variation among clades and was significantly elevated in the "Human French Guiana" and "Sake" groups (Figure S9).

On YPD, CCM was elevated in the "Human French Guiana" clade, but as expected, most of the clade averages hovered around 1 (no CCM).

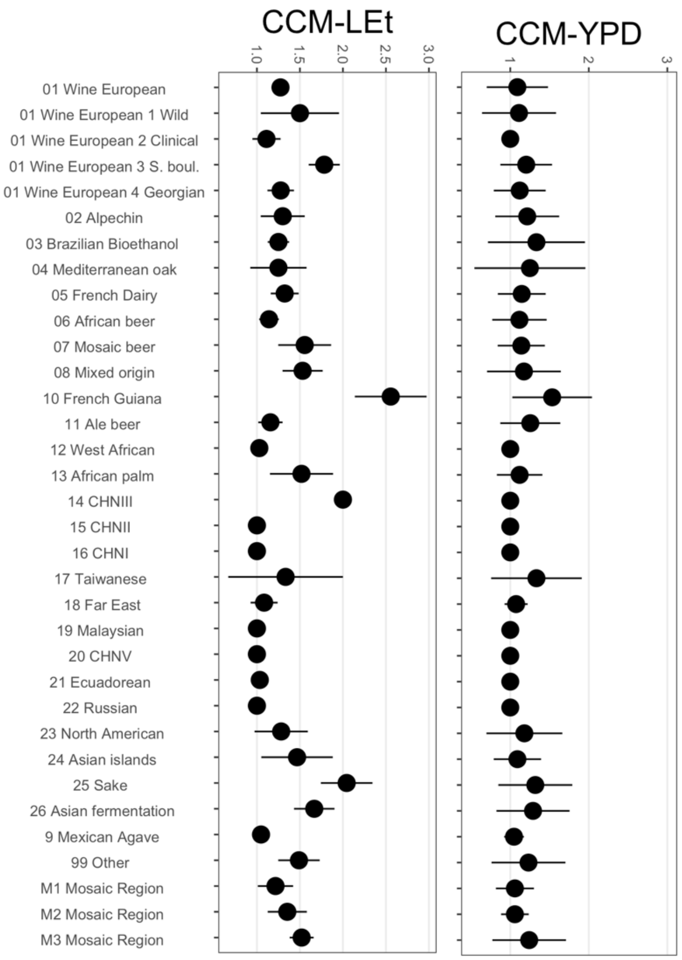


**Figure S9: CCM expression in established clades.**
The plots contain the average expression of all the isolates in a phylogenetic clade (± 2 sem).

*Genetic Analysis*

GWAS was performed on CCM induced each medium. Both genomic control inflation factor ($\lambda_{GC}$) values were close to 1, suggesting that the population structure was correctly stratified. The deviations between the expected and observed -$\log_{10}$(p) values in quantile-quantile plots suggest that there may be strongly associated variants (Figure S10). The list of significant SNPs and associated genes is listed in Table S8.

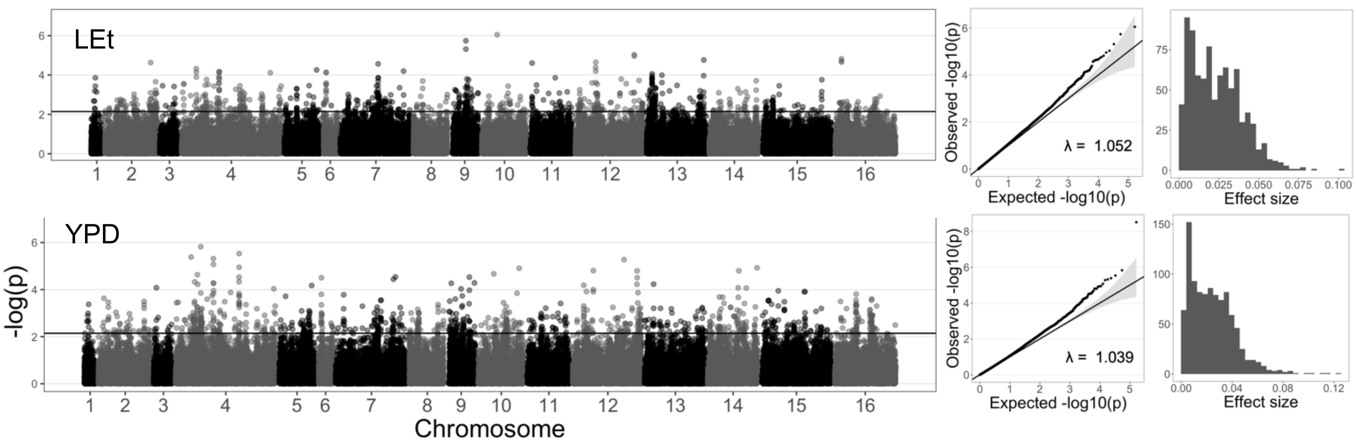


**Figure S10:** Top row- analysis of LEt; bottom row- analysis of YPD. The first column contains the Manhattan plot, which plots the -$\log_{10}$(*p*-values) for the nuclear genomic SNPs; mitochondrial data not included. The second column contains Q-Q plots, which plot the observed *p*-values and the expected *p*-values under a null distribution; each contains a genomic control inflation factor ($\lambda$). The third column contains effect size distributions for the statistically significant SNPs. After normalization, the range of phenotypic values were: LEt -2.9 – 3. and YPD -4.3 – 5.1.
